# A general principle of dendritic constancy – a neuron’s size and shape invariant excitability

**DOI:** 10.1101/787911

**Authors:** Hermann Cuntz, Alexander D Bird, Marcel Beining, Marius Schneider, Laura Mediavilla, Felix Z Hoffmann, Thomas Deller, Peter Jedlicka

## Abstract

Reducing neuronal size results in less cell membrane and therefore lower input conductance. Smaller neurons are thus more excitable as seen in their voltage responses to current injections in the soma. However, the impact of a neuron’s size and shape on its voltage responses to synaptic activation in dendrites is much less understood. Here we use analytical cable theory to predict voltage responses to distributed synaptic inputs and show that these are entirely independent of dendritic length. For a given synaptic density, a neuron’s response depends only on the average dendritic diameter and its intrinsic conductivity. These results remain true for the entire range of possible dendritic morphologies irrespective of any particular arborisation complexity. Also, spiking models result in morphology invariant numbers of action potentials that encode the percentage of active synapses. Interestingly, in contrast to spike rate, spike times do depend on dendrite morphology. In summary, a neuron’s excitability in response to synaptic inputs is not affected by total dendrite length. It rather provides a homeostatic input-output relation that specialised synapse distributions, local non-linearities in the dendrites and synaptic plasticity can modulate. Our work reveals a new fundamental principle of dendritic constancy that has consequences for the overall computation in neural circuits.

**In brief:** We show that realistic neuron models essentially collapse to point neurons when stimulated by randomly distributed inputs instead of by single synapses or current injection in the soma.

**Highlights:**

- A simple equation that predicts voltage in response to distributed synaptic inputs.
- Responses to distributed and clustered inputs are largely independent of dendritic length.
- Spike rates in various Hodgkin Huxley (HH) like or Leaky Integrate-and-Fire (LIF) models are largely independent of morphology.
- Precise spike timing (firing pattern) depends on dendritic morphology.
- NeuroMorpho.Org database-wide analysis of the relation between dendritic morphology and electrophysiology.
- Our equations set precise input-output relations in realistic dendrite models.

## Introduction

Because of their cell-type specific characteristic morphologies, dendritic trees have commonly been assumed to be crucial for a neuron’s intrinsic computations. It has been shown that altering the morphology (Mainen and Sejnowski, 1996; Vetter et al., 2001) or the topology (van Elburg and van Ooyen, 2010; van Ooyen et al., 2002) of neurons while keeping the electrotonic features unchanged has a profound impact on the spiking behaviour of the cell. On the other hand, the morphology of dendrites has been shown to be largely predicted by connectivity rules (Cuntz et al., 2010) rather than by the specific computation that they perform. Also, dendrites were shown to follow general principles that equalise passive (Bird and Cuntz, 2016; Cuntz et al., 2007; Connelly et al., 2016; Jaffe and Carnevale, 1999) and active (Häusser, 2001; Magee, 2000) signal propagation indicating that the blueprints of computation for single neurons might be more stereotypical than previously assumed. In fact, principles of conservative scaling that preserve electrotonic features have been proposed for a number of cell types (Bakken and Stevens, 2011; Bekkers and Stevens, 1990; Cuntz et al., 2013). Similarly, general morphological scaling laws, have been discovered for dendritic arbours of various sizes (Cuntz et al., 2012; Snider et al., 2010; Teeter and Stevens, 2011). However, it remains unclear how electrotonic and morphological scaling principles relate to one another and how their interplay would affect well-known neuronal computations in dendrites (e.g. Branco et al., 2010; Gabbiani et al., 2002; Poirazi et al., 2003b,a; Single and Borst, 1998). Therefore, we study here the dependence of input-output properties of dendrites on their size and shape.

One of the most eminent electrophysiological features of neurons that depend on dendritic shape is the input conductance (Koch et al., 1990; Rall et al., 1967). Smaller cells with smaller input conductances are more excitable for somatic current injections than larger cells because a voltage threshold for spike initiation is reached with a lower input current in accordance with Ohm’s law (Chavlis et al., 2017; Šišková et al., 2014). This relation is true for somatic activation of neurons and its size- and shape-dependence of voltage responses is well understood. However, the corresponding effect of changes in input conductance on voltage responses to distributed synaptic inputs have not been sufficiently studied. Rules identified for current transfer within dendritic arbours (Bird and Cuntz, 2016; Cuntz et al., 2007; London et al., 1999; Rall and Rinzel, 1973; Rinzel and Rall, 1974) have allowed the prediction of responses to individual or a few synaptic inputs (Magee, 2000; Williams and Stuart, 2003). Similar rules should be applicable at the level of richer synaptic input but they have not yet been identified. In this work, we specifically address the question how neuronal firing rate and firing patterns are affected by dendritic size and shape in the case of multiple, distributed synaptic inputs. We show that passive electrotonic principles generally render the synaptically driven excitability of neurons invariable to length for the entire range of existing dendritic trees. Since this dendritic constancy principle supports the stability of neuronal spiking, it may complement other well-established synaptic and intrinsic mechanisms of firing rate homeostasis (Turrigiano and Nelson, 2004).

## Results

The idea behind this work comes from the simple reasoning that while larger neurons are in principle less excitable they also receive more synapses (Figure 1A). The higher input conductance and resulting decreased excitability might therefore compensate for the increase of effective current the neuron receives through its synaptic inputs. In contrast to most traditional theoretical studies on dendritic integration with their focus on somatic activation of the cell or activation with few synapses, we therefore focus here on the voltage responses to distributed synaptic inputs. In the following, we first study these relations analytically in the simple passive cable and subsequently move to passive and then active responses in dendritic trees with their full morphology.

**Fig 1.**
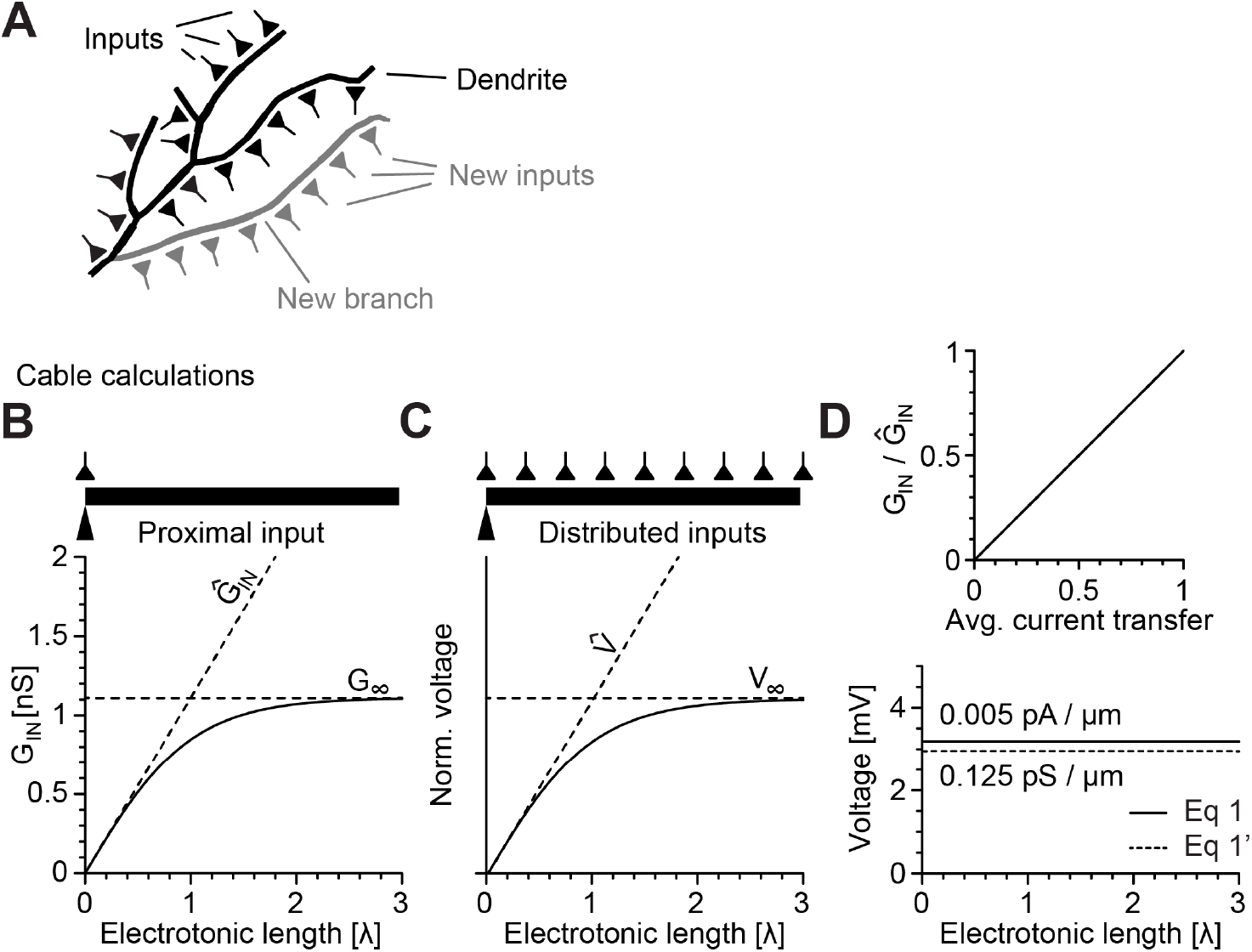
Analytical prediction indicates that responses to distributed synaptic inputs in a cable are independent of cable length. **A**, Sketch illustrating the impact of a new branch on dendrite length and number of synapses. **B**, Input conductance *G*_*IN*_ of cables with constant diameters for a wide range of electrotonic lengths. *G*_*∞*_, the input conductance of a semi-infinite cable and *Ĝ*_*IN*_, the collapsed total membrane conductance are indicated by dashed lines for reference. **C**, Mean steady-state voltage responses to distributed inputs as a function of electrotonic length. *V*_*∞*_, the response to distributed synapses in the semi-infinite cable and 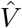, the linear extrapolation of the voltage response at the root and therefore the response in a collapsed cable are indicated by dashed lines. **D**, Bottom panel shows steady-state voltage responses at the proximal end to distributed current injections every *µ*m (straight line, Equation 1) and to synaptic inputs every *µ*m (dashed line, Equation 1’). Top panel shows the average current transfer versus the ratio of *G*_*IN*_ to *Ĝ*_*IN*_ for the cables of varying lengths from **B**. Panels **B—D** were obtained from numerical simulations validating exactly the results of our analytical calculations.

### Analytical calculations for passive cables predict length-invariant responses to distributed synaptic inputs

Experimentally, the input conductance that predicts a neuron’s excitability is most typically obtained from somatic current injection with concurrent somatic voltage measurements using Ohm’s law to relate conductance, current and voltage. The corresponding analytical calculations for a simple dendritic cable are readily available from classical cable theory introduced to neuroscience by Wilfrid Rall. Considerations of current spread in a passive cylinder allow one to predict the input conductance *G*_*IN*_ for any cable of electrotonic length *L* measured in terms of 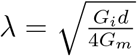, the electrotonic length constant, a distance unit over which the voltage decays to about a third of the proximal voltage (Koch and Segev, 1999; Rushton, 1937). Here, the diameter is *d*, the specific axial conductance is *G*_*i*_ and the specific membrane conductance is *G*_*m*_. For short cables, *G*_*IN*_ increases nearly linearly with *L* as it approximates the collapsed input conductance *Ĝ*_*IN*_ = *G*_*m*_*πdλL* of the cable, the total sum of the membrane leak (Figure 1B). At the other extreme, *G*_*IN*_ at the proximal sealed end in a semi-infinite cable is *G*_*∞*_ = *G*_*m*_*πdλ*, the total conductance of a *λ* length cylinder since *Ĝ*_*IN*_ = *G*_*∞*_*L*. For longer cables of electrotonic length *L*, the input conductance at the proximal end *G*_*IN*_ approaches *G*_*∞*_ asymptotically as *G*_*IN*_ = *G*_*∞*_ tanh(*L*) (Figure 1B). More distal patches of membrane therefore contribute less and less to the total proximal input conductance, setting with *G*_*∞*_ a lower bound for the overall excitability of the cell. In all cases, increasing either the diameter *d*, the specific axial conductance *G*_*i*_ or the specific membrane conductance *G*_*m*_ all increase the input conductance as well as *G*_*∞*_.

As mentioned earlier, apart from their larger input conductance, larger cells also receive more synaptic inputs if one considers constant synaptic density. In the case of very short cables, the increase in input conductance is intuitively perfectly compensated by the increase in number of synapses since both scale linearly with *L*. Interestingly, however, the impact of distal synapses onto voltage at the proximal end diminishes with distance in the very same way as the impact of a distal patch of membrane on the proximal input conductance. The reference values *V*_*∞*_, the average proximal voltage in response to distributed synapses over the dendritic length of a semi-infinite cable, and 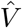, the voltage response of these synapses when they are collapsed to an isopotential piece of membrane, behave similarly to their respective input conductance counterparts *G*_*∞*_ and *Ĝ*_*IN*_ (Figure 1C). It can be shown analytically (see Methods, “Cable equation for responses to distributed inputs”, Equations 3–7) that, along the entire electrotonic length, input conductance and synaptic currents cancel one another precisely. Correspondingly, the average current transfer throughout the cable, i.e. the fraction of injected synaptic current that reaches the proximal cable end, is equal to the ratio of *G*_*IN*_ to *Ĝ*_*IN*_, i.e. the fraction of overall conductance felt at the proximal cable end (Figure 1D, top panel).

The voltage responses to distributed current injections *I*_*dist*_ per unit length are therefore equivalent to the total current injected over the entire metric length *l* = *λL* of the neuron into its collapsed membrane leak, i.e.

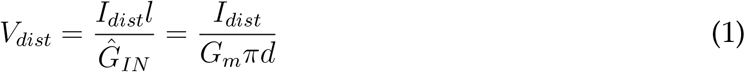

(Figure 1D, bottom panel, straight line). From this particular application of Ohm’s law to dendritic trees, the voltage response to distributed inputs is entirely independent of neuronal cable length while it depends only on the specific conductance per surface membrane *G*_*m*_ and the diameter *d* of the cable as well as *G*_*i*_, since *I*_*dist*_ is defined per unit electrotonic length. This is in stark contrast with voltage responses to proximal “somatic” current injections where 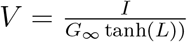, and length impacts the excitability of a neuron by decreasing *V* dramatically when increasing *L*. It is clear, however, that in a realistic setting, excitatory neuronal inputs produce synaptic currents that are distributed over the dendritic tree rather than being somatic. In fact, synaptic currents flow through synaptic conductances that further increase the overall conductance per unit length assuming that synaptic densities are homogeneous and constant. The corresponding voltage responses to distributed synaptic conductances are therefore slightly lower than the ones from current injections. However, also these effects remain independent of total cable length:

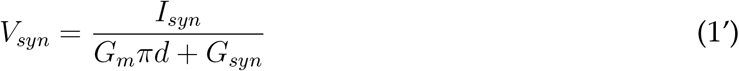

(Figure 1D, dashed line). In conclusion, our analytical calculations and numerical simulations reveal a new electrotonic principle of dendritic constancy ensuring an equal impact of distributed synaptic inputs in a passive cable independent of its length.

### Passive responses encode percentage of active synaptic inputs in a manner that is largely independent of branching topology and dendrite length

Importantly, the principle of dendritic constancy found in the construction of the simple cable with constant diameter can be generalised to branched and tapered neuronal morphologies. We show this at the example of Lobula Plate tangential cells (TCs, *n* = 55) in the blowfly, dentate gyrus granule cells (GCs) in rat (*n* = 43) and mouse (*n* = 8) and cortical pyramidal cells (PCs, *n* = 69) in the monkey with their respective *G*_*m*_ under steady-state distributed inputs (Figure 2). These three datasets were chosen to represent a very leaky large cell (TC), a small and electrotonically compact cell (GC) and the most typical cortical cell (PC) from a range of different species. Normalising the average diameters to the overall average diameter *d* of the respective datasets shows that the steady-state responses are independent of branching patterns and diameter taper (compare larger dark dots with black lines in rightmost panels in Figure 2). In addition, the individual voltage responses of each cell with their original diameters were well predicted by Equation 1 (small dark dots in rightmost panels of Figure 2) with normalised root mean square errors (nRMSE) of 1.3% for TCs, 0.9% for GCs, and 1.2% for PCs.

**Fig 2.**
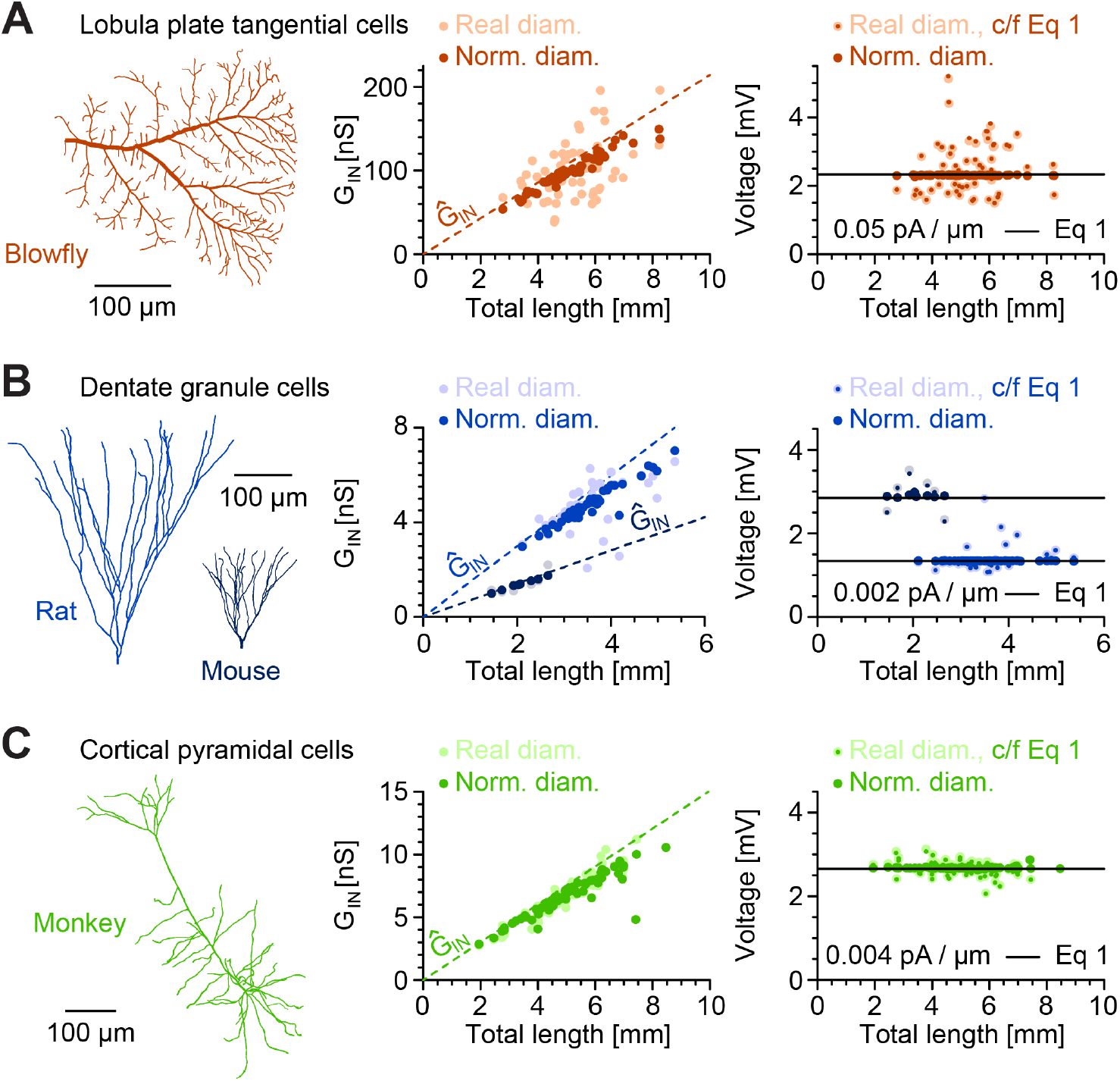
Passive steady-state model responses to distributed synaptic inputs are independent of dendrite length, topology and diameter distribution. Sample morphologies (left), their input conductances (middle panels) and responses to steady-state distributed inputs (rightmost panels) compared with the prediction from Equation 1. **A**, Blowfly Lobula Plate tangential cell (TC) dendrites (red, *n* = 55) with *G*_*m*_ of 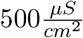; **B**, Dentate gyrus granule cells (GCs) of rat (light blue, *n* = 43) and mouse (dark blue, *n* = 8) with *G*_*m*_ of 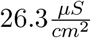 — differences in the species come from different average diameters in the two populations; **C**, Monkey cortical pyramidal cell (PC) dendrites (green, *n* = 69) with *G*_*m*_ of 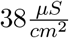. Each dot corresponds to one morphology; lighter dots are original morphologies without diameter normalisation. Large darker dots are results for morphologies with diameters normalised to the average of their respective population; small darker dots are individual predictions from Equation 1 for each non-normalised morphology with its respective average diameter. Straight black lines show predictions from Equation 1 using the average diameter of each population of morphologies and their respective *G*_*m*_. The dashed lines show the collapsed input conductance *Ĝ*_*IN*_ as it increases linearly with the total amount of cable.

Our prediction also accounts for responses to a smaller proportion of activated synapses, i.e. a lower synaptic density, with voltage responses linearly relating with the percentage of active synapses. However, it is important to show what effect a specific, more clustered, distribution of synapses would have on the overall responses in individual neurons. We therefore titrated for any given percentage of active synapses the two most extreme distributions: We compare voltage responses to the activation of a given proportion of the most distal (Figure 3, solid lines) and, respectively, the most proximal (Figure 3, dashed lines) synapses. Even under such clustering of active synapses, neurons seemed to be able to encode the percentage of active synapses with their root voltages both in the steady-state (Figure 3, middle panels) and in dynamic simulations (Figure 3, right panels) following Equation 1’ that includes the synaptic conductances present in these simulations. Importantly, passive somatic voltage responses reflected the percentage of active synaptic inputs independently of morphological complexity and dendrite length (compare Figure 3A with Figures 3B and C).

**Fig 3.**
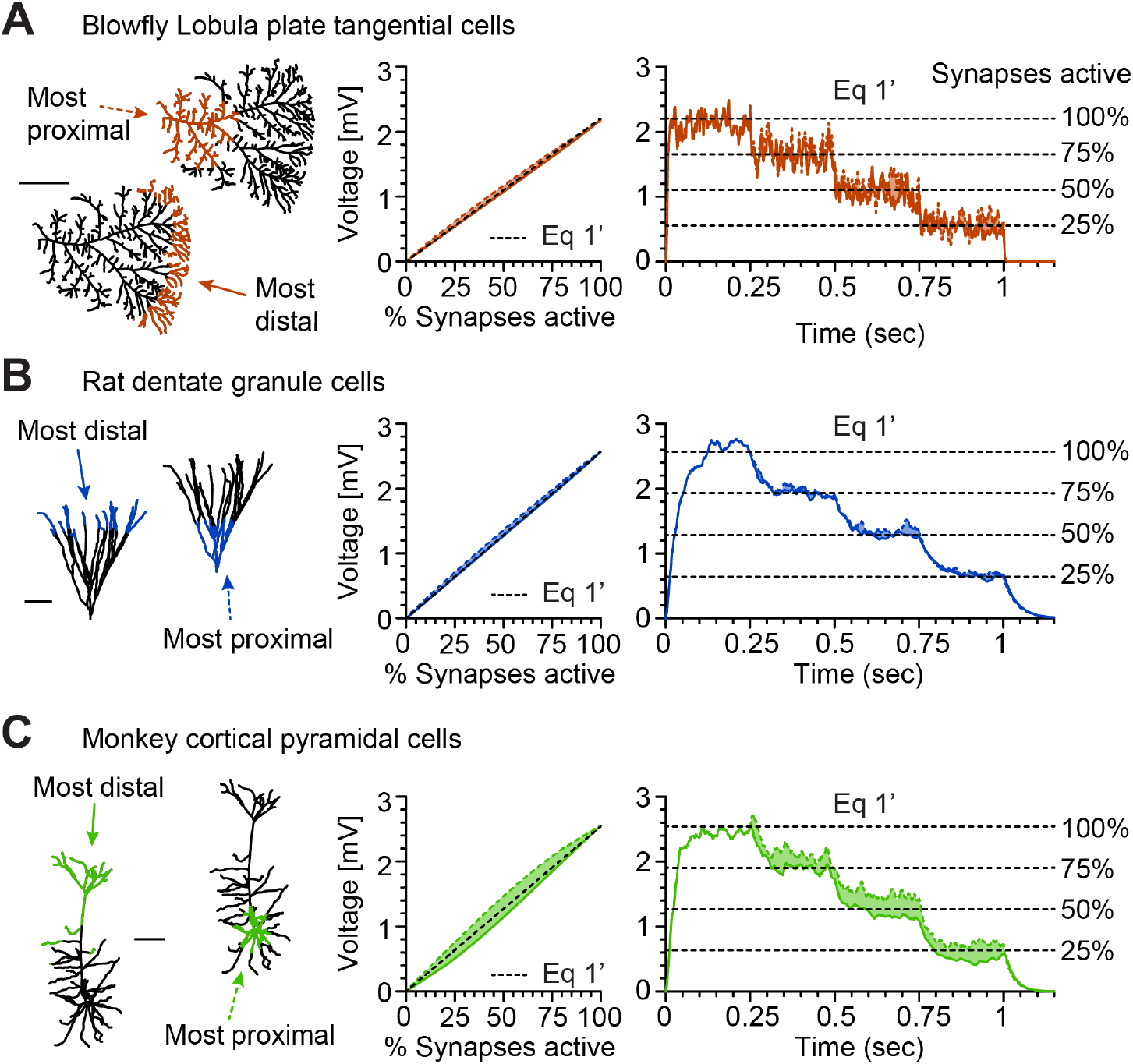
Passive model voltage responses follow relative percentage of active synapses even when these are clustered. Synapse distributions (left), steady-state responses to partial activation of synapses (middle) and responses to sample levels of (100%, 75%, 50%, 25% and 0%) in dynamic simulations (rightmost panels). **A**, **B**, and **C**, each single out one morphology (the one shown on the left) from the populations used in Figure 2 (using the same colour scheme). Dashed coloured lines are the responses to the most proximal synapses while solid coloured lines show the responses to the activation of the most distal synapses. The space in between both responses is shaded. For example, the 25% line means that the 25% most proximal synapses were active (dashed lines) and in a second simulation the 25% most distal synapses were active (solid lines) Black dashed lines are predictions from Equation 1’ that include the synaptic conductance. Scale bars show 100*µm*.

Next, we tested whether the principle of dendritic constancy holds across diverse dendrite 1 branching patterns and sizes in a large number of different cell types. Indeed, our calculation 1 for the steady-state voltage response to distributed inputs in the simple cable yielded good 1 predictions for the wide range of real dendritic morphologies from the July 2016 version of 1 the NeuroMorpho.Org database (Ascoli, 2006). We selected those datasets (223 datasets, 9, 841 1 reconstructions, Table S1) that contained dendritic morphologies with sufficient detail in 1 all three dimensions and with reconstructed diameters (see Methods). Input conductances and steady-state voltage responses to distributed inputs were calculated after normalising the diameters to an average 1*µm* and for generic values of *G*_*m*_ of 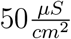 that are typical for cortical pyramidal cells (Figure 4A). We observed here that very large trees exhibited a trend to smaller voltage responses. We found similar results in morphological models for dendritic trees based on minimum spanning trees (Cuntz et al., 2007, 2010, 2012) covering a very large range of possible complexities and overall sizes in synthetic dendrites (Figure S1). Also, the dynamic responses to synaptic stimulation were well predicted for morphologies from NeuroMorpho.Org using Equation 1’ but very small trees showed strong fluctuations because of the small number of synapses there (Figure 4B). Overall, the responses were faithful to our prediction over a range of four orders of magnitude of dendritic length (nRMSE of 5.1% for the steady state responses compared with Equation 1).

**Fig 4.**
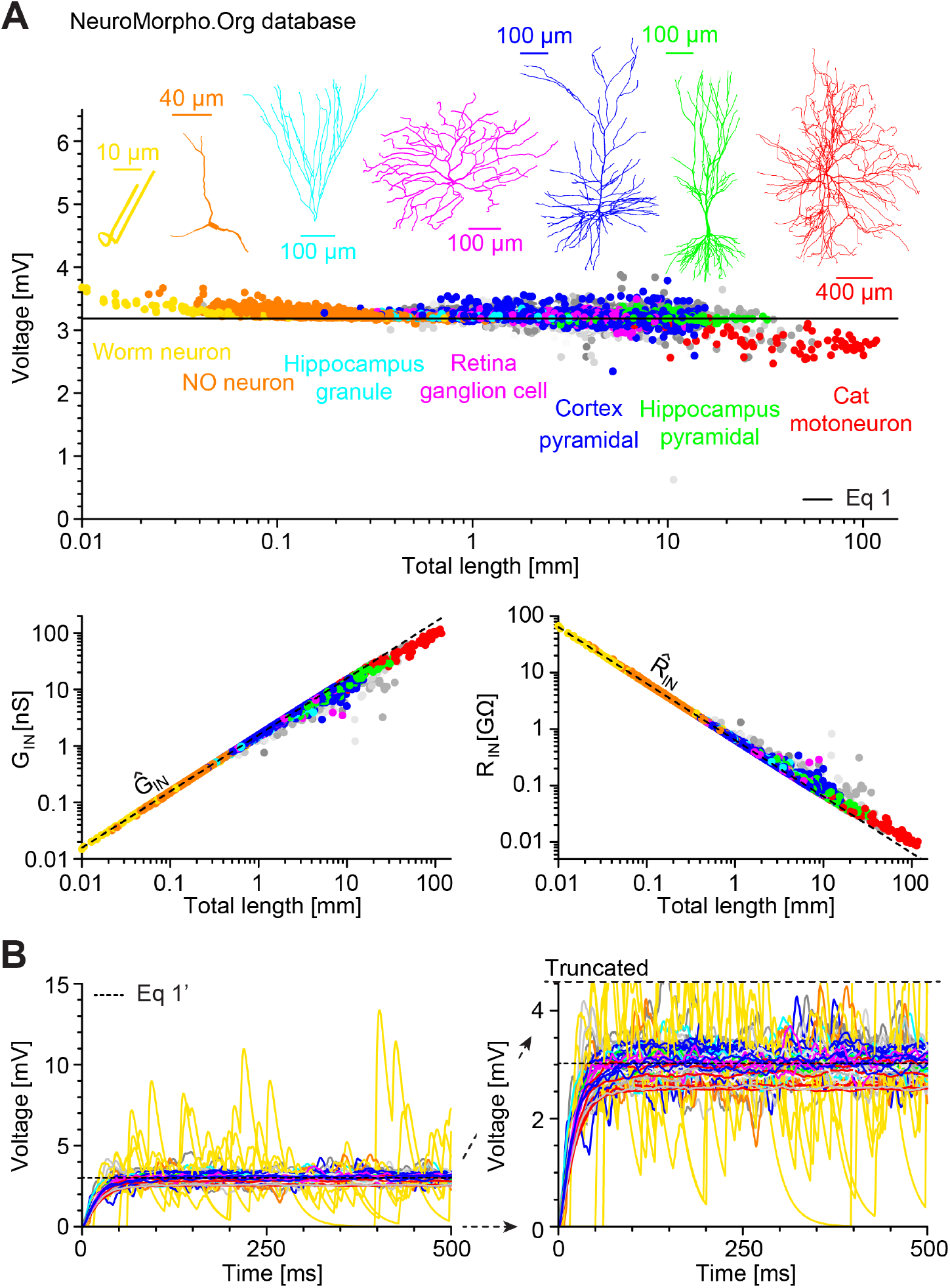
NeuroMorpho.Org database-wide analysis reveals neuronal size and shape invariant passive model responses to distributed inputs. **A**, Voltage responses to distributed inputs with *G*_*m*_ of 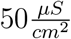 in a large selection of all morphologies from NeuroMorpho.Org (223 datasets, 9, 841 reconstructions, Table S1) after normalisation of average diameters to 1*µm*. Larger consistent subgroups are indicated by colours, representative morphology and label. Unlabelled smaller groups are different shades of grey in the background. The straight line indicates the analytical prediction from Equation 1 for an unbranched cable. Input conductances *G*_*IN*_ (left bottom) and input resistances *R*_*IN*_ (right bottom) are indicated in the same colour code as in the top panel and compared to the case where the overall membrane was collapsed in *Ĝ*_*IN*_ and 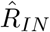 respectively (dashed lines). **B**, Passive dynamic responses and prediction from Equation 1’ (dashed line) in the first morphology of each of the 223 datasets, similarly to the three morphologies in Figure 3, rightmost panels. Since small worm neurons (yellow) exhibited large fluctuations around the mean, this panel is shown at two different scales.

### Spike frequency but not temporal sequence of spikes is independent of model dendrite shape and size in response to distributed synaptic inputs

So far, we have shown a fundamental aspect of passive normalisation of the neural response to synaptic inputs that seems true for all dendritic morphologies. In the following, we used a classical spiking model of cortical pyramidal cells (Mainen et al., 1995; Mainen and Sejnowski, 1996) to test whether our principle of dendritic constancy translates to active spiking neurons. When incorporated in diverse neuronal morphologies from different cell types, this spiking mechanism was previously shown to exhibit strongly varying spiking patterns for somatic current injections (Figure 5A, top row; similar analysis to the original paper using the model #2488 from ModelDB however with normalised average diameters for a better comparison with our predictions, see Methods) (Mainen and Sejnowski, 1996). To quantify the spiking behaviour in four different cell types we plotted firing rates as a function of injected current into the soma (f-I-curves, Figure 5B, top panel). As expected, the spiking frequency increased with decreasing dendrite size rendering smaller cells more excitable. We used interspike interval (ISI) distributions to characterise the temporal structure of the spike trains (bursting vs. non-bursting) in the different morphologies (Figure 5B, bottom panel, ISI distribution refers to single cell firing). L5 pyramidal cells exhibited bursts of three spikes (as indicated by a larger proportion of short ISIs) and L3 pyramidal cells bursts of two spikes (as indicated by two equal peaks in the ISI distributions). Interestingly, when stimulated by distributed synaptic inputs instead of somatic current injections, the differences in the temporal structure of the spiking (bursting vs. non-bursting) remained dependent on the respective morphology with similar ISI distributions as well as coefficients of variation (cv) (Figure 5C, bottom panel). However, the numbers of spikes were equalised and were independent of dendritic tree size irrespective of the frequency of stimulation (Figure 5A, bottom traces, and Figure 5C, top panel). The equalised passive voltage responses predicted by our dendritic constancy (Figure 5C, rightmost panel, bottom traces, subthreshold) were transformed into equal number of spikes in the active model (Figure 5C, rightmost panel, top traces). Taken together, in response to distributed synaptic inputs, spike numbers (frequencies) were independent of morphology while spike times (as reflected in the temporal structure of the spiking in the form of bursting vs. non-bursting) remained affected by morphological properties of dendrites in line with previous observations by Mainen and Sejnowski (1996) for responses to somatic current injections.

**Fig 5.**
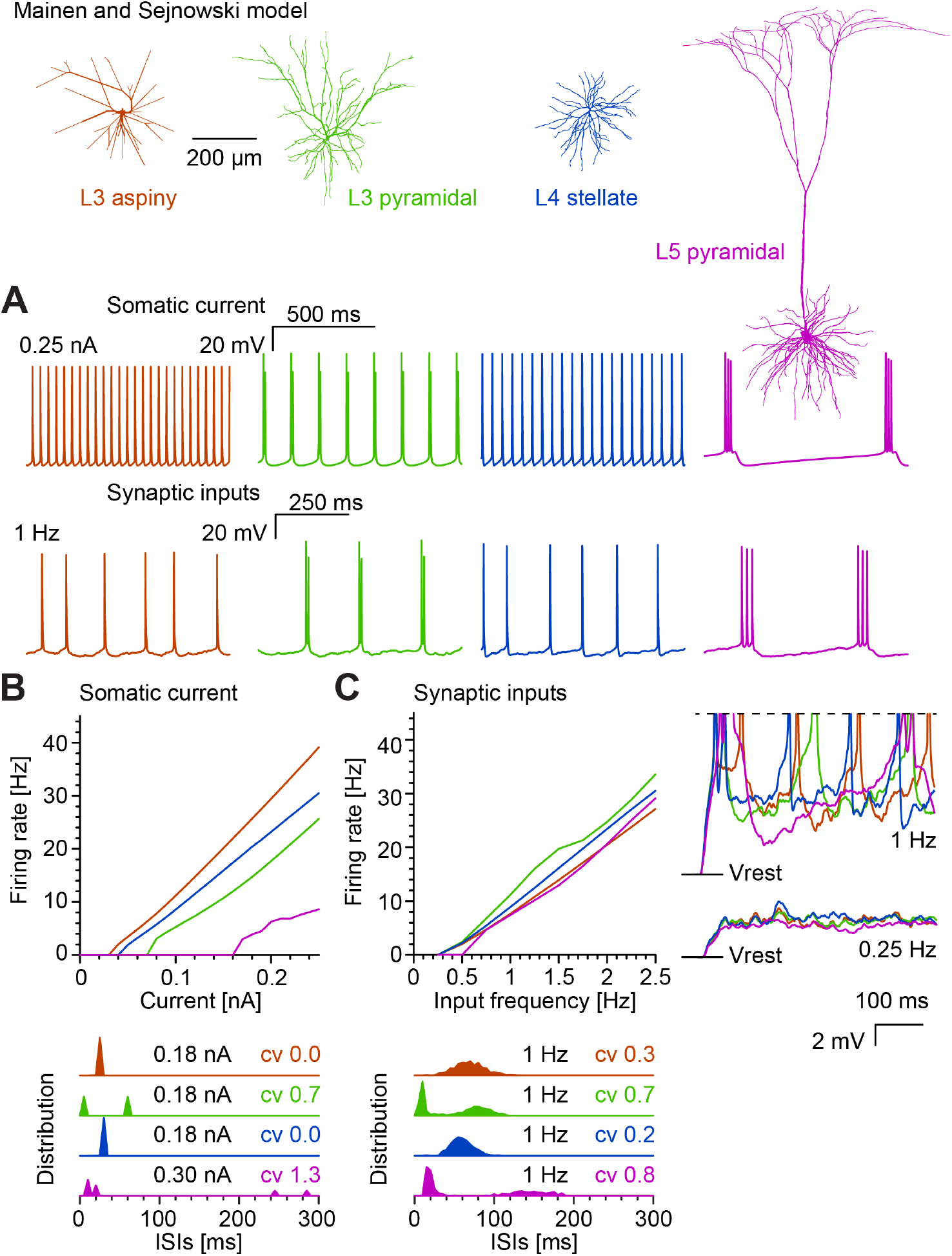
Neuron size- and shape-invariant responses to distributed inputs in spiking neurons. Simulations using the spiking mechanism by Mainen and Sejnowski (1996) in their four sample morphologies of L3 aspiny (dark orange), L3 pyramidal (green), L4 stellate (dark blue), and L5 pyramidal (pink) cells after normalising the diameters. **A**, Sample voltage traces for 0.25*nA* current injections into the soma (top, similar to Figure 1 in the original work; see Methods) and distributed synaptic inputs at 1*Hz* (bottom). **B**, Firing rate vs. current injection in the soma and **C**, responses to distributed synaptic inputs for the same four cases. Sample subthreshold (bottom, 0.25*Hz*) and suprathreshold (top, 1*Hz*; truncated spikes at dashed line) synaptic activation in the four morphologies are shown in the rightmost panels. Interspike interval (ISI) distributions are shown below the respective panels for 40*sec* simulations at indicated current injections and synaptic activation with corresponding coefficients of variation (cv). Note the two peaks in ISI distributions of L3 and L5 pyramidal cells indicative of their bursting. Colours indicate the different morphologies from **A** throughout the figure.

In order to verify that the results in the model by Mainen and Sejnowski were not model-specific, we performed similar simulations in two distinct well-established active models of CA1 pyramidal cells by Jarsky et al. (2005) and Poirazi et al. (2003b). We integrated the corresponding active ion channel models into the set of all good reconstructions of hippocampal pyramidal cell morphologies (*n* = 105) from NeuroMorpho.Org after normalising their diameters to 1*µm* (Figure 6A). The model by Jarsky et al. has previously been used to study the separate effects of inputs from Schaffer collaterals (SC) and the perforant path (PP) (Jarsky et al., 2005). This gave us the opportunity to compare our results for distributed inputs over the entire dendrite with results for inputs that were clustered in a more realistic manner according to their anatomical (layer-specific) origin. In this case we compared stimulating all synapses with 1*Hz* (Figure 6B, black dots) and, separately, only synapses impinging on the basal dendrites (Figure 6B, red dots) or on the distal apical dendrite and tuft region (Figure 6B, cyan dots). Remarkably, in all cases, the firing rates were independent of neuron size. Again, the number of spikes was indicative of the percentage of active synapses scanned in a similar manner to Figure 3 (Figure 6B, rightmost panel). In particular, the corresponding input-output functions were almost identical when measured in two different sample morphologies of radically different total dendritic length (Figure 6B, rightmost panel, compare both sets of solid and dashed lines).

**Fig 6.**
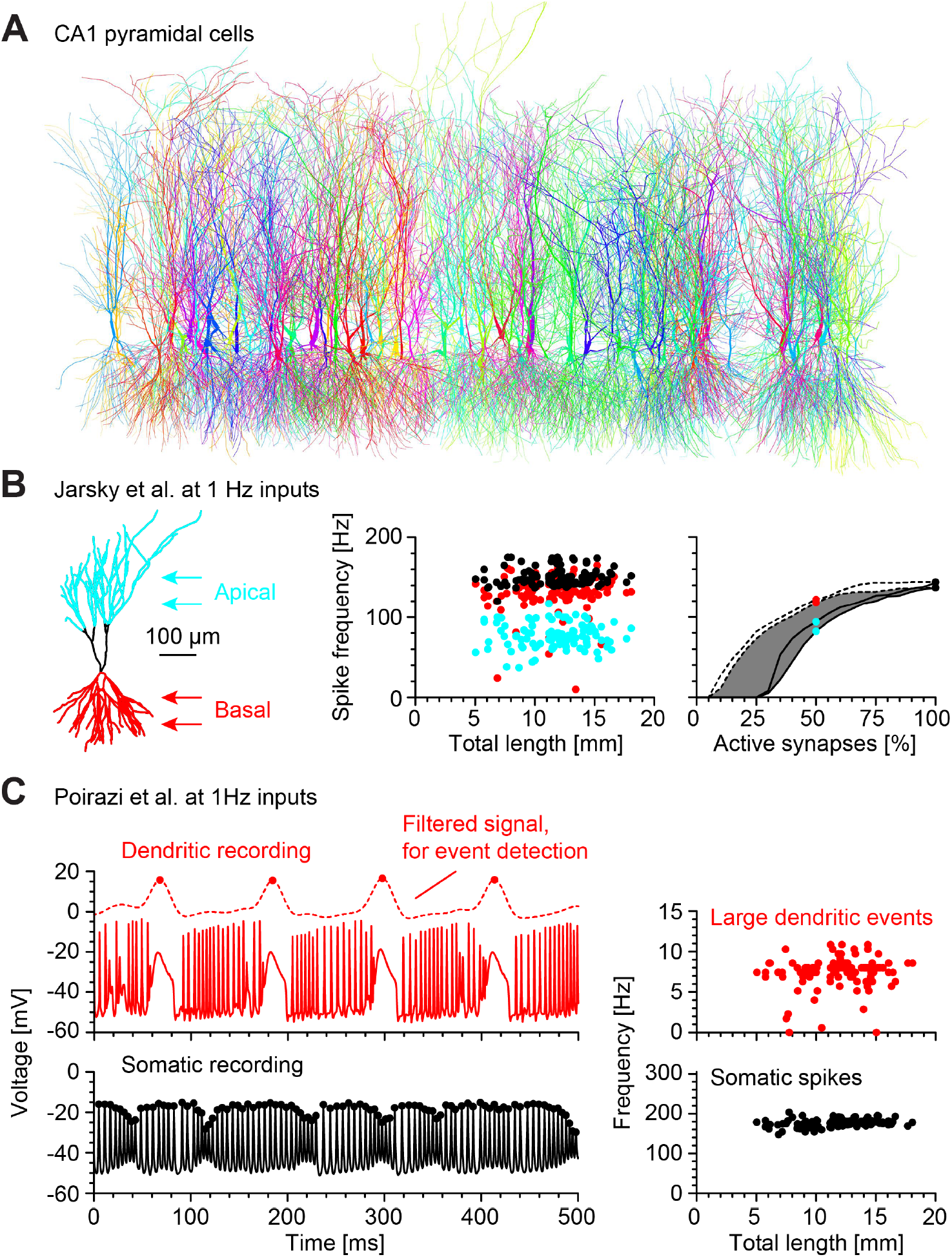
Dendritic constancy in two active models of hippocampal CA1 pyramidal neurons including dendritic spikes and clustered inputs. **A**, All 105 morphologies of rat hippocampal pyramidal cells from NeuroMorpho.Org that passed our manual curation criteria (Colours are random). **B**, CA1 pyramidal cell model by Jarsky et al. (2005) with its responses to distributed synaptic inputs 500*pS*, 1*Hz* in the set of all 105 morphologies. Black dots show spiking responses to activation of all synapses while red (basal) and cyan (apical) show responses to activation of subregions of the dendrite as indicated in the sketch on the left. Rightmost panel shows spike output analysis for selective activation of a subset of all most proximal (dashed line) and most distal synapses (solid line). Since roughly 50% of synapses were active in both the basal and apical stimulations, corresponding values of the curve for proximal (red) and distal (cyan) synapses are highlighted in the rightmost panel as well as values for all synapses (black). The two sets of curves (two dashed, two solid lines, respectively) represent two morphologies of radically different size (10*mm* vs. 15*mm* of total dendritic length). The shaded area highlights the range of responses in clustered synapses for the morphology shown in the inset. **C**, CA1 pyramidal cell model by Poirazi et al. (2003b) driven by distributed synaptic inputs 500*pS*, 1*Hz* in the set of all selected 105 hippocampal pyramidal cell morphologies from NeuroMorpho.Org. Top row, Large dendritic events (red dots) were measured at the branch with highest *y* value respectively and detected after low pass filtering the local voltage signal there with a Gaussian filter with variance of 100*ms* (see dashed line). Bottom row, Somatic events (black dots) as measured directly from the somatic voltage, shown for one sample morphology. Rightmost panels show resulting frequency of somatic (black, bottom) and dendritic (red, top) events with one point each per morphology as a function of total dendritic length.

A second model of CA1 pyramidal cells by Poirazi et al. (2003b) has become archetypal for compartmentalised computations in dendrites. Similarly to the model by Jarsky et al., we incorporated the ion channel models by Poirazi et al. into the NeuroMorpho.Org collection of hippocampal pyramidal cell morphologies and subjected the individual compartmental models to various combinations of distributed synaptic inputs. The model by Poirazi et al. produced large dendritic events that were distinct from the somatic action potentials (Figure 6C, voltage traces red – dendrite vs black – somatic). Intriguingly, both numbers of somatic spikes and large dendritic events were independent of total dendritic length (Figure 6C, left lower and upper panels respectively).

### Spiking reset converts constant membrane voltage into constant spike rates

The results from the active compartmental models pose the question as to why the constant voltages transform into constant numbers of spikes. Such a transformation could be a consequence of each spike essentially shunting and resetting the entire neuron (Hä usser, 2001) while erasing its voltage history. Under these assumptions, a leaky integrate-and-fire (LIF) (Stein, 1965) mechanism coupled to the dendrite could help elucidate the constancy of spike numbers. We chose to implement a LIF that resets the voltage throughout the entire dendritic tree after passing a threshold voltage at the dendrite’s root. Incorporating such a spiking mechanism in the four cell types of Figure 2, yielded an output spiking frequency that was indeed independent of dendritic length (Figure 7A) for any given synaptic input frequency. In fact, the entire input-output (IO) curves were essentially independent of the morphology (Figure 7B). We then derived an analytical solution for the transformation of variable synaptic input activity into firing rate output (see Methods, Equations 10–22). In line with our numerical LIF simulation results (Figure 7A, and B), the mean analytical voltage response to stochastic inputs in a uniform cable was independent of length. The variance of the subthreshold voltage response decayed to a constant for dendrites of total electrotonic length greater than one. Our analytical predictions for the IO relationship (Brunel and Hakim, 1999) are

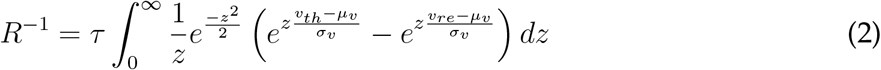

with firing rates *R* for subthreshold voltage mean *µ*_*v*_ and standard deviation *σ*_*v*_ impinging on a membrane with time constant *τ*, firing threshold *v*_*th*_, and voltage reset *v*_*re*_. *R* always converges to constant values for cable lengths longer than the electronic length (see Methods, Equations 10–22, Figure 7B bold black lines). Importantly, *R* is practically independent of dendritic length, if the mean afferent drive is sufficiently strong and so the output firing rate is less dependent on fluctuations (as seen in Figure 7A). In our case in Figure 7B, the specific membrane conductance in the different cell types determined the slope of the IO curves with a very sharp slope in the leaky blowfly TC that in reality produces spikelets and shallower curves in the other cell types with lower conductivity through the membrane. Interestingly, the percentage of active synapses was encoded in the number of spikes (Figure 7C) in analogy to the voltages in Figure 3 (compare with IO curves in Figure 7B). Again, this was true even for clustered synapses (plots show most distal synapses as solid lines vs. most proximal synapses as dashed lines).

**Fig 7.**
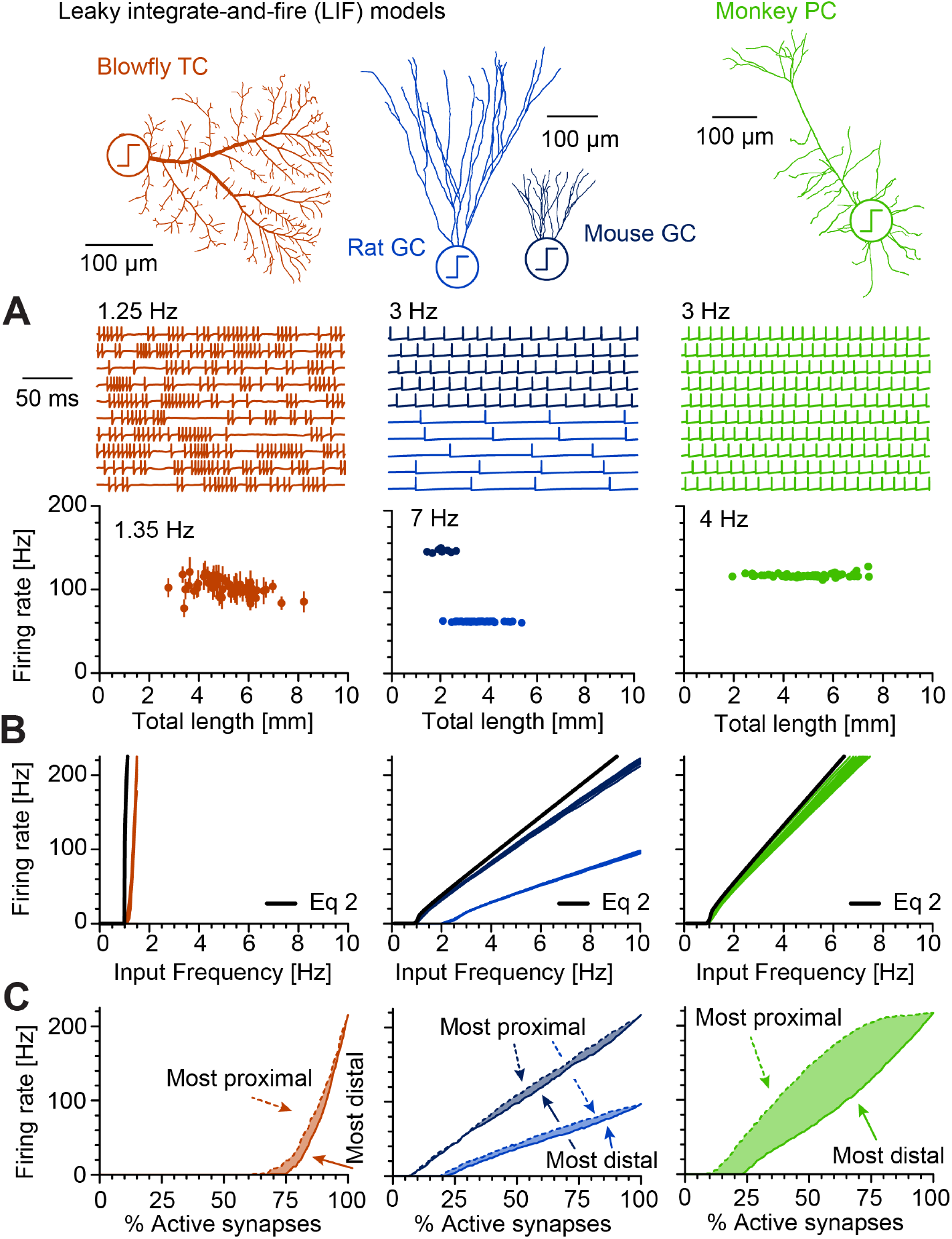
Spiking responses to distributed inputs in leaky integrate-and-fire (LIF) neurons with realistic dendritic morphologies as the source of leak indicate that voltage reset throughout the entire dendritic tree contributes to spike rate constancy. LIF mechanism in its simplest form (without adding soma or axon, the LIF resets the voltage in the entire dendrite after spiking) introduced into the dendritic root of the cell types from Figure 2 with the same colours and membrane properties. **A**, Top row, Sample spike trains for 10 different TC, 5 rat GC, 5 mouse GC and 10 monkey PC dendritic morphologies are shown underneath the corresponding morphologies. Bottom row, Firing rates are shown with error bars (standard deviation, invisible in GCs and PCs) for one selected input frequency for each cell type are shown as a function of length. **B**, Input-Output (IO, Frequency of synaptic activation vs. spiking frequency) plots for all available morphologies for TCs (left), mouse and rat GCs (middle) and monkey PCs (right). Respective cable calculations from Equation 2 that are independent of length are shown as bold black lines. **C**, Responses to selective activation of the most proximal (dashed lines) and most distal (solid lines) synapses in the dendrite. The areas between these two extreme scenarios for clustered synapses are shaded.

Similarly, incorporating the LIF mechanism into the dendritic morphologies of the Neuro-Morpho.Org database (with a uniform specific membrane conductance, see Figure 4) yielded invariant IO curves over a very large range of morphologies. Only the IO curves from tiny worm neurons (yellow) and very large spinal cord motoneurons (red) deviated from the remaining curves (Figure 8A, LIF model, compare also these results with analytical predictions from Equation 2). In the same morphologies, we also showed spike number invariance using an adaptive exponential LIF (AdExpLIF) (Brette and Gerstner, 2005) for two specific temporal patterns of spikes typically seen in compartmental models, a bursting mode and spiking mode with spike frequency adaptation (Figure 8B and C, AdExpLIF model). Overall, LIF based spiking models were consistent with the dendritic constancy of passive voltage responses to distributed synaptic inputs transforming into constant spike numbers that were independent of dendritic length or shape.

**Fig 8.**
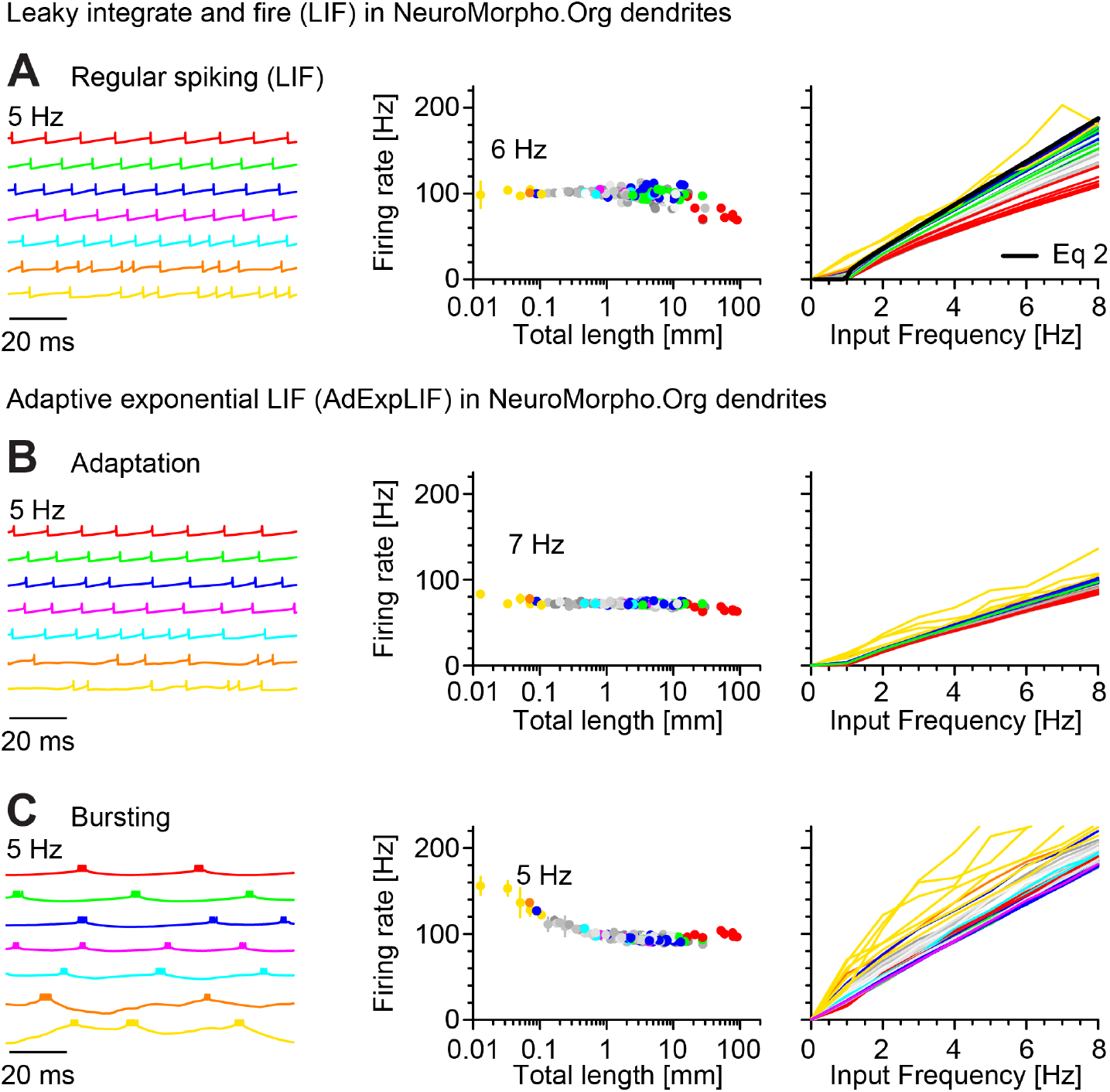
NeuroMorpho.Org database-wide spiking responses to distributed inputs in leaky integrate-and-fire (LIF) and adaptive exponential LIF (AdExpLIF) neurons are independent of dendritic size and shape. Similar panels as in Figure 7A and B but for all morphologies from Figure 4 with the respective membrane properties used there: Sample voltage traces in sample morphologies for all seven morphological categories (left panels), firing rates in response to a given input frequency (middle panels, input frequency is indicated), and IO curves (right panels). Colours as in Figure 4 in increasing dendrite size: Yellow (worm neurons); orange (nitergic neurons); cyan (hippocampal granule cells); pink (retinal ganglion cells); blue (cortical pyramidal cells); green (hippocampal pyramidal cells); red (cat motoneurons). **A**, Regular leaky integrate-and-fire (LIF) spiking mechanism incorporated in all dendritic morphologies. Similarly to Figure 7, calculations from Equation 2 are shown as a bold black line. **B**, Adaptive exponential LIF (AdExpLIF) that includes an additional channel for spike frequency adaptation and results in less regular spiking. **C**, Long time constant in the adaptation channel of the AdExpLIF for testing our dendritic constancy theory under extreme burst firing.

## Discussion

In this work, we used analytical methods to demonstrate a general principle of dendritic constancy regarding the voltage and spiking responses to distributed synaptic inputs. Synaptic inputs effectively encounter an apparent input conductance in the soma (i.e. the transfer conductance) corresponding to the collapsed membrane leak of the entire dendrite onto the soma. As a consequence, more synaptic currents in larger cells are precisely compensated by the additional dendritic leak. Our dendritic version of Ohm’s law (Equation 1 and Equation 1’ as well as Equation 2 for spikes with variable inputs) is independent of morphological features and spiking mechanisms and predicts isoelectrotonic behaviour for anatomically distinct dendritic trees shaped by species-specific scaling (Beining et al., 2017; Cuntz et al., 2013), developmental expansion (Mckay and Turner, 2005) or neurodegenerative shrinkage (Platschek et al., 2016). Finally, our simulations in a classical model by Mainen and Sejnowski (1996), as well as other established spiking models (Jarsky et al., 2005; Poirazi et al., 2003b) and LIF models showed that synaptic stimulation in different dendritic trees leads to similar responses in terms of firing rates but not patterns (spike times). This was true for uniformly activated synapses but also to a large degree for clustered synapses, so much so, that the somatic firing rate allowed for an approximate estimation of the percentage of active synapses independent of their dendritic location. Taken together, our analytical and numerical results imply that the principle of voltage and spike rate constancy is general since it holds in all (branched and unbranched) dendritic arbours activated by distributed synaptic conductances.

### Limitations for dendritic constancy

What are the assumptions and limitations of our computational analysis? First, dendritic constancy will be affected by a number of dendritic features. The voltage responses do depend on the specific membrane conductances and average dendritic diameters. While average dendritic diameters do not seem to vary much in the NeuroMorpho.Org database (Figure S2), *G*_*m*_ values are known to vary between cell types (Borst and Haag, 1996) (Figures 2, 3 and 7) and also within cell types (e.g. Garden et al., 2008). In addition, the somatic membrane leak affects the dendritic constancy results when somata are very large compared to the overall dendritic membrane if their diameters are not normalised together with the dendrites (i.e. if they do not scale with dendrite length) and if they do not receive synaptic inputs (Figures S2 and S3).

Second, in our analysis, we assumed uniform kinetics, conductances and reversal potentials of synaptic inputs. In this respect, it would be interesting to further explore models with variations in distance-dependent synaptic properties (Häusser, 2001; Magee and Cook, 2000) as well as democratising effects on distal synapses (London and Segev, 2001; Rudolph and Destexhe, 2003). Here, we show for a distance-dependent linear gradient of maximal synaptic conductance (increasing with dendritic path length from the root) that dendritic constancy is preserved in the hippocampal CA1 pyramidal cell model by Jarsky et al. and in the NeuroMorpho.Org morphologies (Figures S4A and B). Furthermore, our analytical solutions predict that dendritic constancy applies also to inhibitory synapses with negative reversal potentials (same models, Figures S5A and B). However, the effects of layer-specific somatic or dendritic inhibition, introducing local or distant shunts (Gidon and Segev, 2012) and their interaction with active channels (see below) need to be studied in detail.

Third, it also remains unclear how dendritic non-linearities such as dendritic NMDA or calcium spikes would operate in the context of our dendritic constancy principle. Such non-linear computations would include dendritic integration features known to play a role depending on relative locations of synaptic inputs (Branco et al., 2010; Cuntz et al., 2003; Poirazi et al., 2003b; Polsky et al., 2004). Surprisingly, our active cortical and hippocampal models exhibited dendritic constancy (without requiring any specific tuning) of spike numbers and even in the numbers of dendritic active events in the model by Poirazi et al. (Figures 6C). However, many further aspects related to dendritic voltage-dependent channels remain to be explored. For instance, potassium channels and H-channels are capable of affecting local synaptic potentials as well as backpropagating spikes (Chen et al., 2006; Magee, 1999). It would be intriguing to test how these channels shape dendritic constancy for synchronous or asynchronous, clustered or distributed synaptic inputs.

Fourth, it remains to be determined how dendritic constancy might interact with recently described homeostatic plasticity of the axon initial segment (AIS) in the form of activity-dependent changes in its location and length and in the distribution of its ion channels (Adachi et al., 2015; Kuba, 2012). Decreased or increased synaptic activity can induce homeostatic lengthening or shortening of the AIS with a compensatory increase or decrease in neuronal excitability respectively (Evans et al., 2015; Kuba et al., 2010). However, the effect of AIS length or location on excitability is more complex and depends on neuronal size (Gulledge and Bravo, 2016). Therefore, further computational and experimental analyses are needed to better understand the link between the neuron’s size and shape invariant excitability that we describe here and AIS plasticity.

Of particular interest is our observation that spike times rather than spike numbers remained affected by morphological properties of dendrites in a similar manner to responses to somatic current injections (Mainen and Sejnowski, 1996). Our simulations revealed that in the case of synaptic stimulation, both dendritic constancy as well as variability in firing patterns were maintained at the same time in active models of four different reconstructed cortical cell types. Although two cortical cell models displayed regular firing and the other two bursting, all of them generated similar firing rates. The somatic bursting behaviour has been previously explained as a consequence of delayed dendritic depolarisations and subsequent return currents from dendrites, arising due to two key factors: (1) moderate coupling resistance between somatic and dendritic regions in combination with (2) separated distributions of fast and slow active channels in soma and dendrites (Mainen and Sejnowski, 1996). Our synaptically driven simulations confirmed and extended these analyses by showing that the electrotonic mechanisms of dendritic constancy are able to normalise spike numbers in cells with different dendritic sizes and shapes without disrupting the active burst generating mechanisms. It is tempting to speculate that dendritic constancy could support homogeneous spike-rate coding across different morphologies while at the same time allowing for cell-type specific spike-time coding. In other words, dendritic constancy may facilitate neuronal computations by maintaining stable firing rates while keeping variability of spike patterns (Denève and Machens, 2016; Denève et al., 2017; Gjorgjieva et al., 2016).

### Clinical relevance of dendritic constancy

The dendritic constancy principle could be of clinical relevance. Changes in dendritic size and shape are hallmarks of many neurological disorders, including chronic stress (Conrad et al., 2017), stroke (Brown et al., 2010; Qin et al., 2014) and neurodegeneration (Šišková et al., 2014; Spires and Hyman, 2004). Whereas dendritic atrophy caused by direct damage to a neuron is considered part of the disease process (Šišková et al., 2014), dendritic remodelling occurring in disconnected brain areas, i.e. network damage, is most likely homeostatic and restorative in nature. For example, the perforant pathway to the dentate gyrus degenerates in aged humans and in Alzheimer’s disease (Leal and Yassa, 2013; Yassa et al., 2010). As a consequence, the target neurons of this pathway – dentate granule cells – retract their dendrites. This dendritic retraction is caused by denervation and not by the disease itself (Einstein et al., 1994). Experimental animal data have shown that such denervated and retracted granule cells (Vuksic et al., 2011) eventually achieve synapse densities on their dendrites comparable to pre-denervation levels (Steward et al., 1988). In that case, the input conductance as well as the number of synapses with additional unit length would likely cancel each other out. The dendritic tree has fine-tuned itself to achieve firing rate homeostasis (Platschek et al., 2016, 2017). We show here that this feature is not specific to dentate granule cells and that synaptic excitability of neurons is size-invariant for all dendritic trees due to a general electrotonic principle. Thus, transneuronal dendritic remodelling appears to play a homeostatic role in maintaining information throughput in a partially damaged network.

### Practical consequences for computational modelling and input-output computation of neurons

Equations 1 and 1’ allow for quantitative predictions of voltage responses to distributed synaptic inputs. This can be helpful for tuning large-scale morphologically realistic compartmental models (e.g. Markram et al., 2015) because by setting synaptic conductances to a specific value, it is possible to achieve a target voltage (and corresponding spike numbers from Equation 2). Thus, dendritic constancy simplifies the estimation of a neuron’s behaviour within a network. For instance, for a given synaptic conductance, the frequency of synaptic activation required to reach a particular membrane voltage can be computed. From the perspective of network computations, the principle of dendritic constancy can be viewed as a mechanism for preserving stable neuronal activity in the circuit (as done in Figure 7). Intuitively, adding new synapses to a spiking network model would create more spikes. Even one additional spike can dramatically alter network dynamics (London et al., 2010). However, dendritic constancy is one possible mechanism to prevent this from happening, because the cell’s number of output spikes depends on the relative number of active synapses and not on their absolute number. This means that increasing the number of synapses while adjusting the morphology accordingly would effectively not change the total number of spikes in the network.

In summary, our principle of dendritic constancy serves as an equalising homeostatic mechanism on which dendritic non-linearities and synaptic plasticity can operate (London and Häusser, 2005; Turrigiano, 2017). It creates a passive backbone for the conservation of excitability converting a neuron to a reliable size- and synapse number-independent “summing point” within the network (Segev and London, 2000) but at the same time, it allows for more complex computations with active dendrites (Schmidt-Hieber and Nolan, 2017). Because dendritic constancy is based on basic electrotonic properties, it applies to all neurons receiving distributed excitatory or inhibitory inputs. This simple and universal principle has previously been overlooked because most studies focused on neuronal firing activated by somatic current injections or by few synaptic inputs instead of distributed synaptic stimulation. Dendritic constancy becomes apparent after leaving the “somatocentric” and embracing the “synaptocentric” view of a neuron’s input-output transformation.

## Acknowledgments

We would like to thank S. Jagannath, S. Platschek and S. Rozada for performing preliminary analyses and A. Castro, F. Effenberger and M. Schölvinck for useful discussions and comments on the manuscript. The work was supported by BMBF (01GQ1406 – Bernstein Award 2013 to H.C.; OGEAM 031L0109B to T.D.), Deutsche Forschungsgemeinschaft (CRC 1080 to T.D.), University Medical Center Giessen and Marburg (UKGM; to P.J.), LOEWE CePTER – Center for Personalized Translational Epilepsy Research (to P.J. and T.D.) and F.Z.H. was supported by the International Max Planck Research School (IMPRS) for Neural Circuits in Frankfurt. The authors declare to have no competing financial interests.

## Author contributions

H.C., A.D.B, M.B, M.S., L.M., F.Z.H., T.D. and P.J. conceived the study and wrote the paper. H.C., M.B., M.S., L.M. and F.Z.H. performed the numerical simulations and A.D.B. performed the analytical calculations.

## Materials and methods

### Data and algorithm availability

All passive electrotonic and leaky integrate-and-fire (LIF) simulations were done in Matlab (Mathworks Inc, 2015b, 2017b and 2018b) using our own open-source software package, the TREES toolbox (Cuntz et al., 2010) (www.treestoolbox.org, Interim version). TREES toolbox functions are marked in italic and end with a *tree* suffix throughout the Methods section. Active compartmental model simulations were done in NEURON (Carnevale and Hines, 2004) using our new software T2N to communicate with the TREES toolbox in Matlab (Beining et al., 2017). All results were further analysed in Matlab. All dendritic morphologies were downloaded from www.NeuroMorpho.Org (Ascoli, 2006) in July 2016. The active model for the spiking mechanism by Mainen and Sejnowski (1996) for Figure 5 used model #2488 from ModelDB (Hines et al., 2004). The LIF and adaptive exponential leaky integrate-and-fire (AdExpLIF) models (Brette and Gerstner, 2005) using realistic dendritic leak in Figures 7 and 8 were implemented in Matlab. All new functions (*cgin_tree*, *LIF_tree*, *LIF_FR_tree*, *AdExpLIF_tree*) will be made available as part of the TREES toolbox on publication at www.treestoolbox.org via Github. The code and data for all figures will be made available at https://zenodo.org/ on publication. The code was tested on various operating systems. Individual methods are detailed in the following but can best be appreciated in the actual Matlab scripts.

### Cable equation for responses to distributed inputs

The voltage response at distance *x* along a closed cable of length *l* due to current of magnitude *I*_*app*_ injected at the root (Rall, 1959, 1962) is

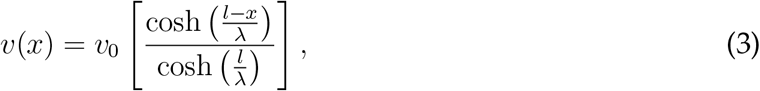

where *λ* is the electrotonic length constant and *v*_0_ is the voltage at the root:

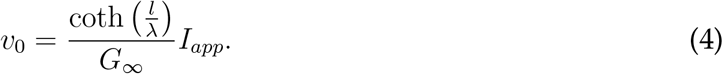

As transfer resistance is symmetric in dendrites (Koch and Segev, 1999; Rall et al., 1967; Rushton, 1937) this also gives the voltage *v*_*x*_(0) at the root due to current injection at a distance *x*

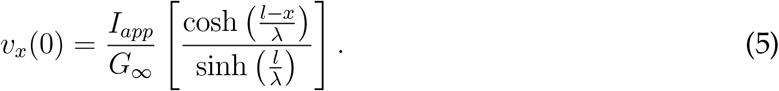

To find the total voltage *V* for currents injected along the entire cylinder, we require the integral over all synaptic sites

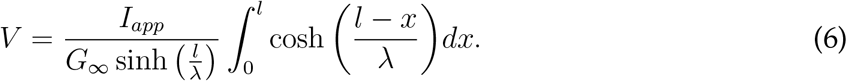

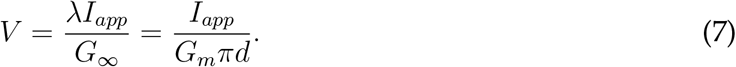

### Morphologies for passive electrotonic simulations

Simple cables (12.5*µm*—12.5*mm* length in 12.5*µm* steps) of constant 1*µm* diameter (Figures 1B–D) or various dendritic morphologies (Figures 2–4) were resampled to constant 1*µm* internode resolution (using `resample_tree`). Individual datasets used in combination with specific membrane properties were from blowfly Lobula Plate tangential cells (TCs, *n* = 55) (Cuntz et al., 2008) (Figures 2A and 3A, 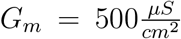), rat (Rihn and Claiborne, 1990) (*n* = 43) and mouse (Schmidt-Hieber et al., 2007) (*n* = 8) dentate gyrus granule cells (GCs, Figures 2B and 3B, 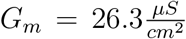) and monkey cortical pyramidal cells (Luebke et al., 2015; Coskren et al., 2015) (PCs, *n* = 69, Figures 2C and 3C, 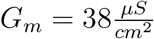). The three datasets were also used in the context of leaky integrate-and-fire (LIF) spiking models in Figures 7. Dendrite morphologies from the entire NeuroMorpho.Org database were used for Figures 4 and 8 by manual curation of all existing archives to select those with sufficient diameter profiles, sufficient depth information in *z*, sufficiently high-quality reconstructions and no sudden jumps in *z* (selection, 223 datasets, 9, 841 reconstructions, see Table S1). The selected archives were sorted by cell types into the following categories in decreasing order of total cable length: Spinal cord motoneurons (red), hippocampal pyramidal cells (green), neocortical pyramidal cells (blue), retinal ganglion cells (pink), hippocampal granule cells (cyan), nitrergic neurons (orange), *C. elegans* neurons (yellow), and other (different shades of grey per dataset). These categories were chosen as representatives for the possible scales of dendrites rather than because they corresponded to consistent cell types. All dendrite morphologies were normalised to a given average diameter (to the average diameters in their specific archives for Figures 2 and 7 and to 1*µm* for Figures 4 and 8).

### Passive steady-state measures for dendritic morphologies

The collapsed input conductance was measured by summing up the leak conductance over the entire membrane surface of the dendrite using the function `surf_tree`. The resulting calculation is made available in the new TREES toolbox function `cgin_tree`. Remaining electrotonic features are all readily available from the electrotonic signature (`sse_tree`) as introduced previously (Cuntz et al., 2010). Briefly, all membrane and axial conductances are arranged according to the tree’s adjacency matrix and the current transfer between all nodes is obtained by taking the inverse of the resulting conductance matrix. Local input resistances are then found on the diagonal of this electrotonic signature since current there is injected in the same node as the voltage is measured. Voltage responses to distributed inputs are simply the sum over the column or row of the electrotonic signature since the matrix is symmetric and the system is linear. While this method simulates steady-state distributed current injections, synaptic conductances associated with batteries according to their specific reversal potentials can be simulated instead (using the `syn_tree` function). The passive results were obtained in their purest form in dendrites without the associated somata and axons. Only for Figures S3 was the effect of somata explored in detail (see below).

### Passive dynamic responses to distributed synaptic conductances

Synaptic inputs were simulated as a Poisson process inducing synaptic conductances at a given frequency per synapse. The dynamics of the conductance trace was given by the form 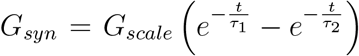 with a rise time constant of *τ*_2_ = 0.5*ms* and a decay time constant of *τ*_1_ = 2.5*ms*. *G*_*scale*_ was set using Equation 1 to ensure that the integral over time of the synaptic conductance profile produced the same voltage as the steady state cases (compare Figure 2 rightmost panels with rightmost panels in Figure 3) at an input frequency of 5*Hz* per synapse. Our novel TREES toolbox function *LIF tree* was used without a voltage threshold for spiking in the case of the passive dynamic responses. *LIF tree* injects distributed synapses into the conductance matrix that defines the dendritic tree in a time-resolved dynamic manner and produces local voltage responses throughout the dendrite. In Figures 3 and 4 the voltage time courses at the dendritic root were plotted for a subset of morphologies (the ones shown in Figure 3 and the first morphology in each of the 223 datasets in Figure 4) for better clarity.

### Effect of soma size on dendritic constancy — analytical treatment

Consider an electrotonically compact soma of radius *R* attached to a dendritic cable of length *l* and radius *r*. The intrinsic properties are given by the specific conductance of the intracellular medium *G*_*i*_ and membrane conductance *G*_*m*_. The soma has a leak conductance of *G*_*s*_(*R*) = 4*πR*^2^*G*_*m*_. The voltage along the cable due to a current injection of magnitude *I*_*app*_ at the soma is given by

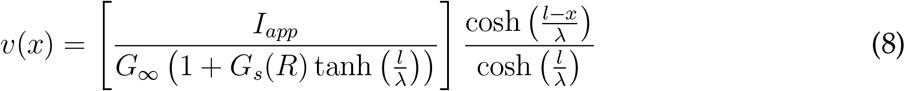

for 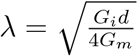, the electrotonic length constant of the cable, and 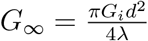, the semi-infinite conductance. Note that this derivation relies on a self-consistent description of the root voltage *v*_0_ due to the current flowing into the dendrite. Due to the symmetry of transfer resistance, this is also the voltage induced at the soma by current injection at a site *x*. Consider the total somatic response *V*_*Tot*_ to distributed synaptic currents:

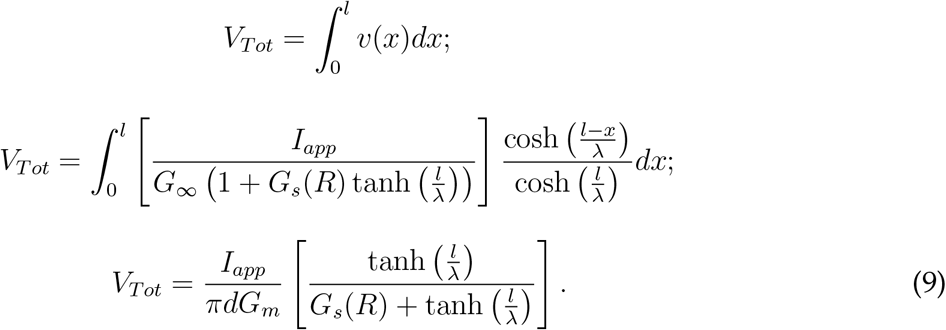

It can be seen that the term in brackets determines the deviation from dendritic constancy. A small value of *G*_*s*_(*R*) is key as tanh is bounded by one. Figures S3 plots the relationship between somatic radius *R* and dendritic constancy for different electrotonic lengths 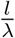 and the relationship between the electrotonic length 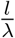 and dendritic constancy for somatic radii *R*.

### Spiking model by Mainen and Sejnowski

We used our new tool T2N (Beining et al., 2017) to port the existing model for the spiking mechanism by Mainen and Sejnowski (1996) #2488 from ModelDB (Hines et al., 2004) to our TREES toolbox package in Matlab. In T2N, calculating spike frequency vs. current injections or vs. synaptic input frequencies using different dendritic morphologies becomes easy to implement. The required simulations are distributed automatically on the available computing cores and the entire toolset from the TREES toolbox becomes available to better edit and analyse dendritic trees and the resulting simulation variables. The code is available but, briefly, the simulations ran 40*sec* with a time step of 0.05*ms* and a pre-run for 200*ms*. The initial voltage was set to *−*70*mV*, which was a close match for resting voltages for the four different morphologies. The voltage was calculated every 50*µm* and a current injection electrode was inserted into the root or synapse point processes into every node (separated by 1*µm*). Morphologies from the original model were translated into TREES toolbox and resampled to 1*µm* internode distances. The dendritic diameters were normalised to 1*µm* and soma with axon divided into axon hillock, initial segment, nodes of Ranvier and myelinated segments were added as in the original model with the respective ion channel conductances. Implicit spines were modelled according to the original model for the current injections but even the L3 aspiny cell was implemented as spiny in all cases for better comparison. Responses to distributed synaptic inputs were modelled with `Exp2Syn` point processes with rise time constant of *τ*_2_ = 0.2*ms* and decay time constant of *τ*_1_ = 2.5*ms* driven by `NetStim` point processes in artificial point neurons under Poisson process conditions (noise 1) and following a given input frequency. The random seeds for the `NetStim` process were set to be independent for different synapses.

### CA1 pyramidal cell spiking models

Electrotonic compartmentalisation and location dependent ion channel distributions allow for separate non-linear integration of inputs in different regions of dendrites. In order to check how these conditions affect our results, we studied two models of CA1 pyramidal cells that are known to produce dendritic spikes. The dendritic arborisation of pyramidal cells follows a laminar structure that generally reflects the different main excitatory afferents impinging on their dendrites from different brain regions. This distinctive structural organisation is also manifested in the way the electrotonic properties and active channels are distributed. Therefore, it was necessary to define how non-uniform channel distributions scale in the different morphologies.

### Jarsky et al. 2005 model

We ported the model by Jarsky et al. (2005) to T2N in a similar manner as with the model by Mainen and Sejnowski. This model includes four active conductances: a voltage-gated Na^+^ conductance, a delayed rectifier K^+^ conductance, a proximal A-type K^+^ conductance, and a distal A-type K^+^ conductance with a higher half-inactivation voltage. These conductances were distributed as a function of path distance from the soma. The Na^+^ and the delayed rectifier K^+^ conductance were modelled following a uniform distribution, the weak excitability version of the model by Jarsky. The A-type K^+^ current was modelled with the experimentally reported six-fold increase in conductance along the apical dendrites resulting in variable slopes of the linear increase between soma and tuft in different morphologies. The apical dendrites were divided with borders along the apical trunk to contain 3.14% (proximal apical), 36.27% (medial apical), 68.90% (distal) and 100% (tuft) of the total apical length respectively. These divisions occurred at path distances of around 100*µm*, 300*µm* and 500*µm*.

### Poirazi et al. 2003 model

The model by Poirazi et al. (2003b) was also ported to T2N, and similarly adapted to apply to different pyramidal cell morphologies. The model consists of a wide variety of active and passive membrane mechanisms (see the online supplement in Poirazi et al., 2003b), including 17 types of ion channels, most of them non-uniformly distributed along the somato-dendritic axis. The apical trunk stems were divided according to laminar depth from soma to stratum lacunosum-moleculare (*>* 68.90% from the total apical dendrite length, similarly as in the model by Jarsky) and the ion channel distributions were rescaled accordingly. The apical trunk dendrites that bifurcate within the stratum radiatum giving rise to two or more main apical dendrites were also considered as the apical trunk region. Similarly to the original Poirazi model, a peritrunk region was defined as the first 50*µm* in path length from every oblique branch that extended away from the apical trunk. The remaining apical branches were considered as the apical region with a further distinction of more distal dendrites, located beyond a laminar depth away from the soma of 300*µm* (distal apical) and 350*µm* (tuft). The passive parameters and channel densities were similar to the Poirazi model, except for axial conductances being distributed uniformly and the leak reversal potential being fixed to *−*70*mV* rendering slightly different resting potentials for each cell morphology.

### Integrate-and-fire spiking model with passive dendrite leak

Dynamic LIF spiking responses for all morphologies in Figures 2–4 were obtained using the `LIF tree` function in a similar way as for passive dynamic responses (see above). In the case of the LIF responses, synaptic conductances were set using Equation 1’ to reach *−*60*mV* at the dendrite root when activated at 1*Hz*. By then setting the voltage threshold of the LIF mechanism in the dendrite root to *−*60*mV* we ensured that spiking started around 1*Hz* input frequency (Figure 7B). Spikes were generated throughout the dendrite when the threshold was reached at the dendritic root, resetting the voltage everywhere to *−*70*mV*. Morphologies from NeuroMorpho.Org were used in their pure dendritic form (without soma or axon) and after normalising dendritic diameters for each population.

### Adaptive exponential integrate-and-fire spiking model with passive dendrite leak

Since the simple LIF is generally not able to reproduce the variety of temporal firing patterns that occur in real neurons we extended it by an adaptation current in combination with an exponential activation term (Brette and Gerstner, 2005), while preserving passive parameters. This also allowed us to test yet another spiking mechanism for our theory of dendritic constancy. Instead of a fixed threshold for spike initiation, action potentials in the adaptive exponential leaky integrate-and-fire (AdExpLIF) are generated through a positive, exponential feedback in the voltage of the dendritic root *V*_*root*_, given by the differential equation 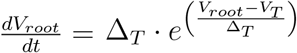. By setting the slope factor to ∆_*T*_ = 2*mV* and the threshold to *V*_*T*_ = *−*60*mV* we made sure that spiking started around 1*Hz* input frequency similar to our LIF simulations. The exponential activation term in the soma makes precise processing of fast fluctuating inputs during synaptic bombardment possible (Fourcaud-Trocmeé et al., 2003), as spike initiation is not instantaneous in contrast to the LIF. The upswing of the potential beyond *−*60*mV* grows rapidly to infinity, which is why the exact numerical threshold for a voltage reset has almost no influence on spike timing and was set to *V*_*thres*_ = 10*mV* in all simulations. Altering the parameters of spike initiation had no effects on the constancy of spike numbers with respect to morphology (*>* 50*mV*). The adaptation current *w* acted as a negative feedback on the voltage in each segment of the dendritic tree and was given by:

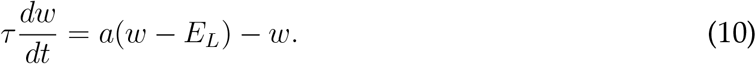

Once the dendritic root reached *V*_*thres*_, the voltage in each node was reset to *V*_*reset*_ = *−*70*mV* in the case of spike frequency adaptation. Increasing the reset voltage to *−*60*mV* induced bursting. After a spike was triggered, the variable *w* was increased by an amount *b* in all segments, which was *b* = *−*60*fA* in the adaptation and bursting neuron model. Depending on *b*, the bursting neuron elicited several spikes in a short period of time until *w* counterbalanced the exponential activation term, resulting in a longer ISI in between bursts. In case of the bursting model, the time constant was set to 30*ms*. Increasing the time constant to *τ* = 100*ms* in combination with a high value of *b* resulted in spike frequency adaptation.

### Stochastic inputs: Subthreshold voltage moments

Consider a sealed dendrite of physical length *l* with electrotonic length constant *λ*. The voltage time course at the proximal end due to a single brief injection of current of magnitude *a* at electrotonic position 0 ≤ *x* ≤ *l* is given by

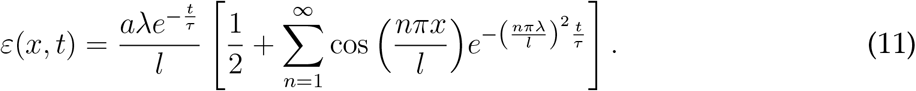

This is plotted in Figure S6A (dashed lines). Given that synapses are uniformly distributed over [0, *l*], the expected (ensemble) value of *ε* at a given time *t*, 〈(*ε*(*t*)〉, is given by

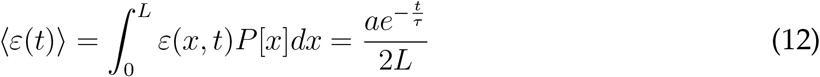

where we have written 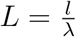 for the electrotonic length. The voltage above rest at the soma, neglecting for the moment a threshold-rest mechanism, is given by a sum of independent synaptic inputs

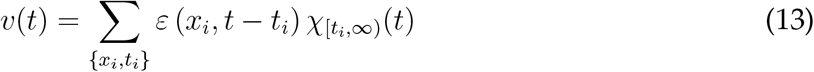

where the times *t*_*i*_ are given by a Poisson process of rate *r*_*λ*_*l*, the locations *x*_*i*_ are uniformly distributed along the dendrite, and *χ*_[*ti,∞*)_(*t*) is the indicator function of the interval [*t*_*i*_, ∞).

The subthreshold steady-state mean voltage above rest 〈*v*〉 can be found by taking expectations

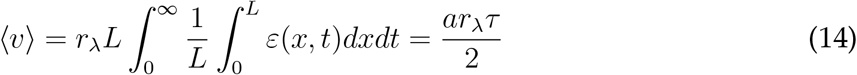

This is independent of *L*. Similarly, the subthreshold variance in *v* can be written as

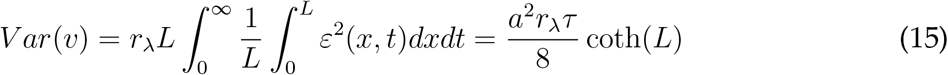

where the coth(*L*) term approaches 1 in the limit of large *L* (Figure S6B, dashed lines).

### Stochastic inputs: Subthreshold voltage moments for synaptic currents

The above calculations give the voltage impulse response at the soma. If a synapse has its own time course *ζ*(*t*) (with *ζ*(*t*) = 0 for *t <* 0), then the somatic voltage above rest is given instead by

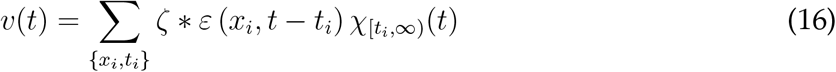

where 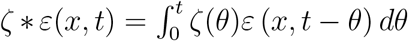 represents convolution in time. A typical synaptic filter is modelled as a difference of exponentials with timescales *τ*_*f*_ and *τ*_*s*_ such that

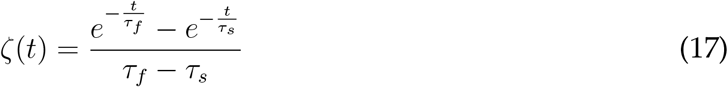

Note that each term in the series form of *ε*(*x, t*) can be written as 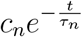 for some coefficient *c*_*n*_ and timescale *τ*_*n*_ as defined above (with *c*_*n*_ typically dependent on input location *x*). Then each such term convolves with *ζ*(*t*) to give

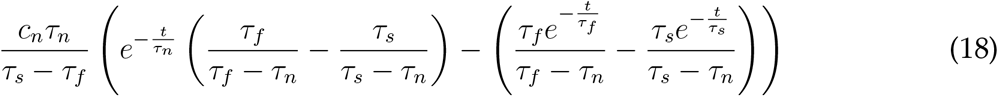

if *τ*_*f*_, *τ*_*s*_ ≠ *τ*_*n*_. In the case that one of *τ*_*f*_ = *τ*_*n*_ or *τ*_*s*_ = *τ*_*n*_ (without loss of generality let *τ*_*f*_ = *τ*_*n*_) the form is instead

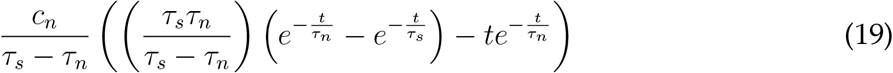

with an additional synaptic filter alongside the dendritic filter given by Equation 11, the difference in somatic voltage responses to proximal and distal inputs is reduced even for synapses that are fast compared to the membrane time constant (Figure S6A).

The subthreshold mean is unchanged from the instantaneous case as 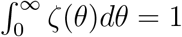, and the subthreshold variance can be computed by squaring the above terms, integrating *t* from 0 to *∞* and *x* from 0 to *l*, and summing the infinite series. The result is cumbersome to write in full, but can be plotted in Figure S6B. The variance is lower in the case of the synaptic filter compared to instantaneous current injection.

### Stochastic inputs: Subthreshold characteristic functions and firing rate approximation

The firing rate can in principle be calculated exactly from the expected time for the stochastic process (Equation 13) to first reach the firing threshold *v*_*th*_ from the voltage reset *v*_*re*_. Given a uniform initial voltage *v*_0_ (which decays with timescale *τ*), the random variable 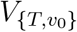 describes the voltage *T* seconds later. The characteristic function 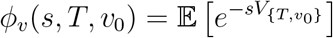 of 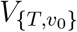 is given by (Rice, 1944)

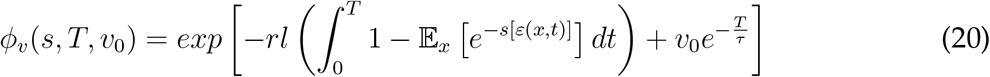

where the expectation 𝔼_*x*_ is over synaptic locations *x*. This can be inverted to give the probability distribution *f*_*v*_ of 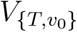. An additional integral transform over *T*, 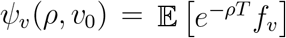, allows the moment generating function *M*_*FP*_(*t*) of the first-passage time density to be written as (Siegert, 1951)

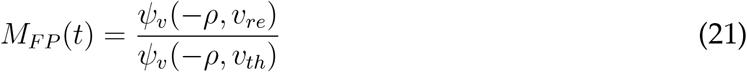

The mean first-passage time, and hence the output firing rate, could then be extracted from 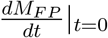.

In practice, the above procedure is numerically sensitive and the following approximation is robust to the high cumulative input rates typically seen across an entire dendritic tree. Taking the subthreshold voltage mean *µ*_*v*_ and standard deviation *σ*_*v*_ allows the firing rate *R* to be accurately approximated (Alijani and Richardson, 2011) using the equation from Brunel and Hakim (1999)

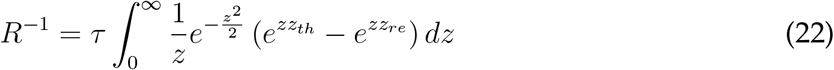

where 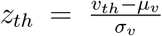 and 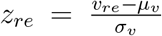. This is plotted as a function of dendrite length in Figure S6B and as a function of input firing rate in Figure S6C.

Combining the above equations, the output firing rate *R* can be written, in the case of instantaneous synapses, in terms of intrinsic quantities as

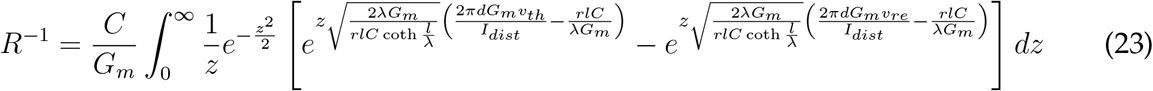

where, as before, *C* is the specific capacitance, *G*_*m*_ is the membrane conductivity, *l* is the dendrite length, *d* is the average diameter, 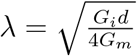 is the electrotonic length, *G*_*i*_ is the axial conductivity, and *I*_*dist*_ is the current induced by a single synapse. Additionally, *r* is the rate of synaptic activation per *µm*, and *v*_*re*_ and *v*_*th*_ are the reset and threshold voltages respectively.

In the case of filtered synapses, there is not a compact form for *R* and Equation 22 is used directly with the subthreshold mean and variance as derived above. The code to calculate *R* analytically can be found in the function `LIF_FR_tree` for synaptically filtered current injections.

## Supporting information

**Fig S1.**
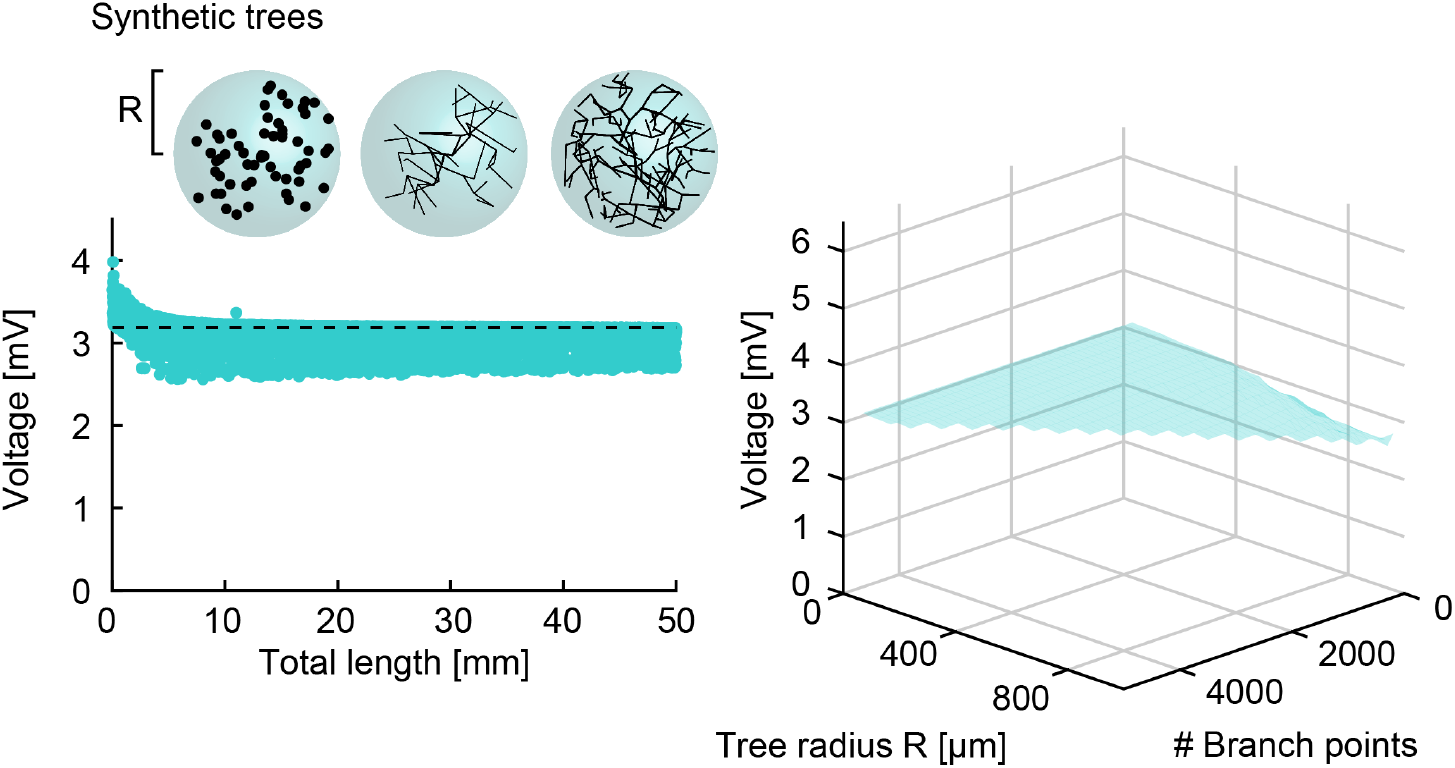
Steady-state passive responses to distributed inputs in synthetic dendrites are independent of dendrite length and shape. Similar analysis as in Figure 4 but for 10, 000 synthetic dendritic trees obtained using extended minimum spanning trees that reproduce many features of real dendrites (Cuntz et al., 2010). These cover a wide range of tree complexities as well as overall sizes (see Methods).

**Fig S2.**
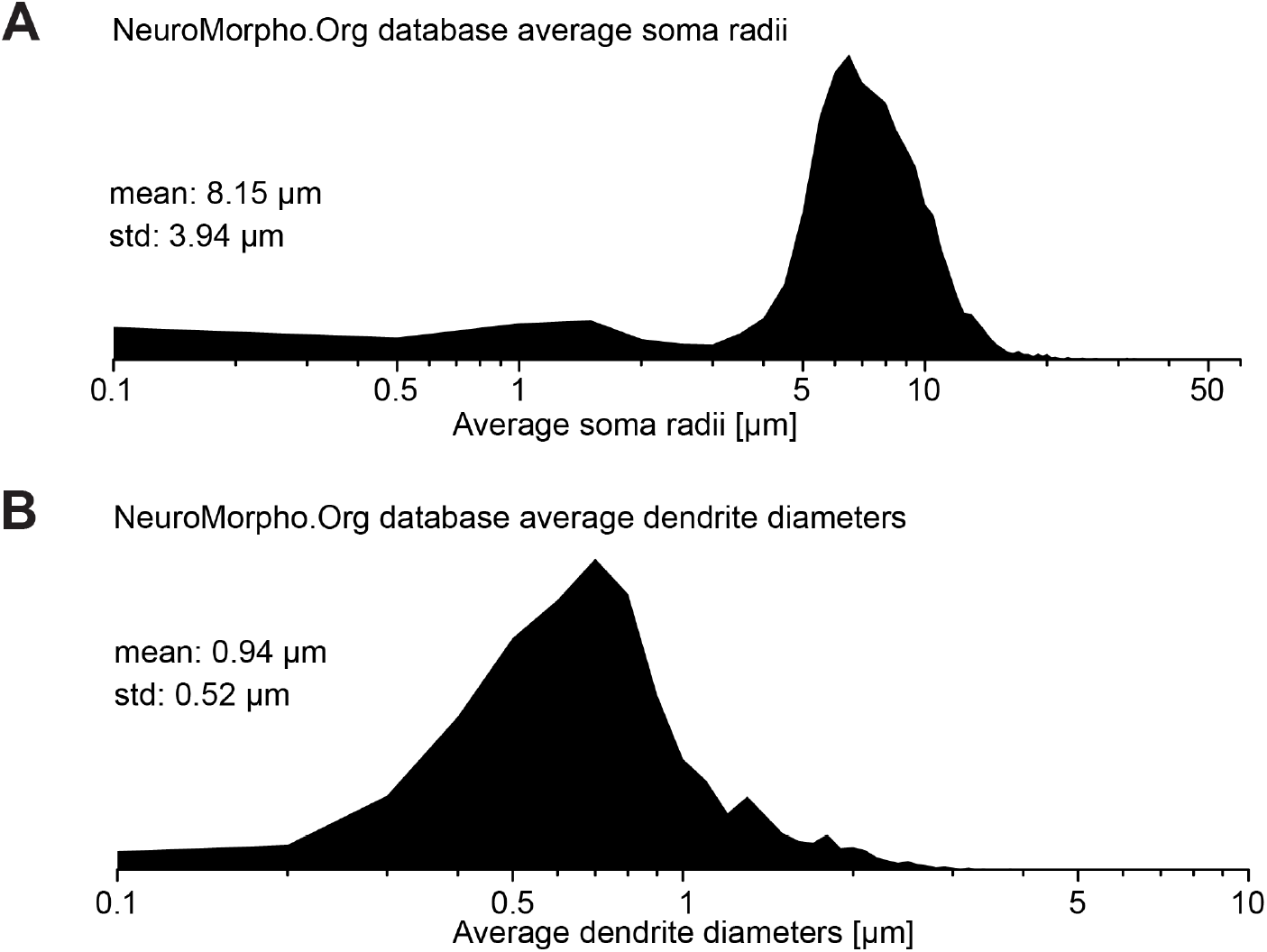
Distributions of diameters in somata and dendrites of the NeuroMorpho.Org database. **A**, Distribution of soma radii in NeuroMorpho.Org. Not every cell had well-reconstructed somata explaining the tail of very small radii. High-quality soma reconstructions were not an inclusion criterion for this study that focuses on dendritic trees in the main text. **B**, Distribution of average dendrite diameters after resampling to 1*µm* internode distances to weigh each location in the dendritic tree homogeneously.

**Fig S3.**
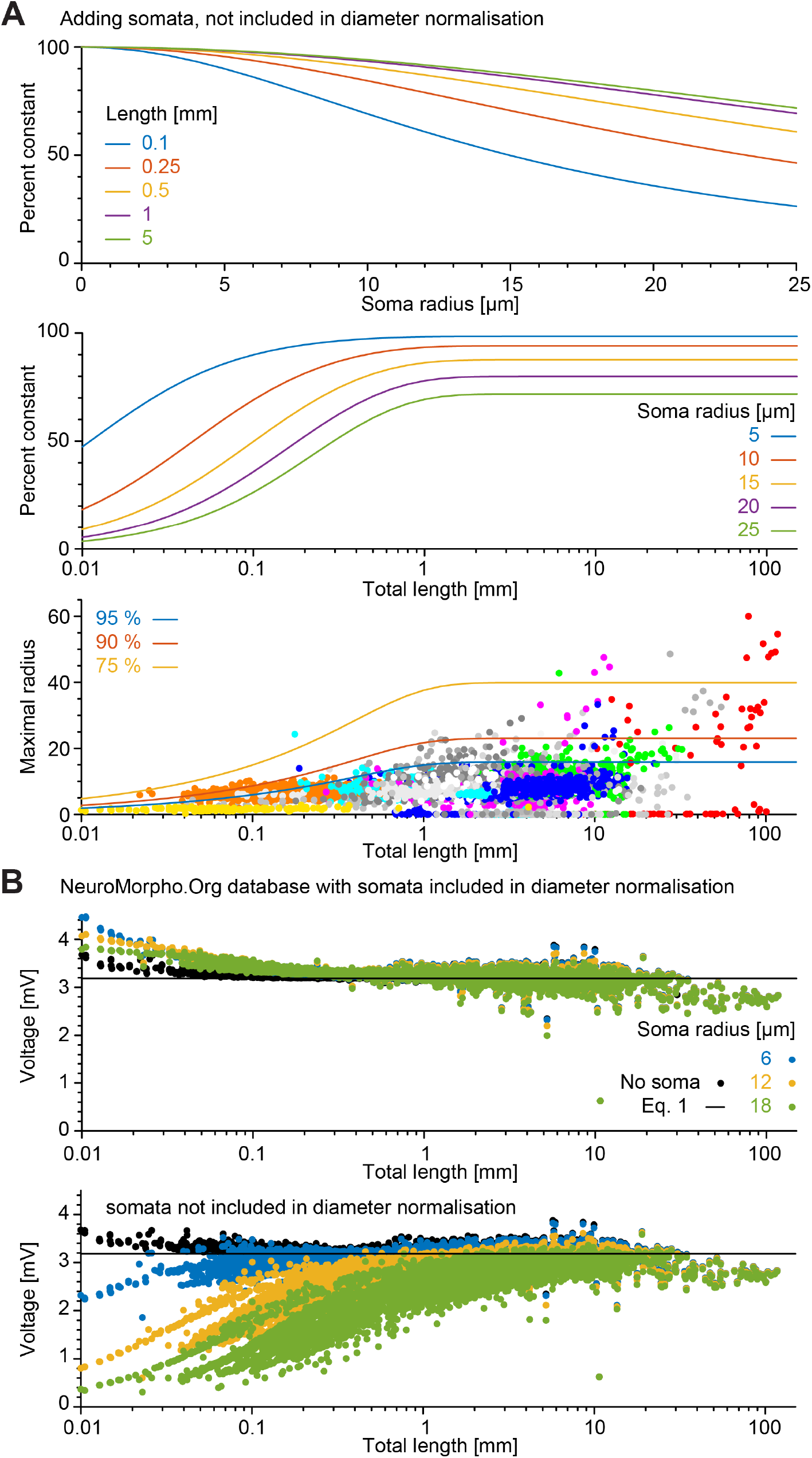
Effect of soma size on dendritic constancy. **A**, Analytical calculations of relative deviation from dendritic constancy (compared to 100%) as a function of somatic radius for different lengths of dendrite (top panel) and as a function of length for different somatic radii (bottom panels). Used cables had 1*µm* diameter, specific membrane conductance 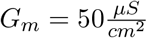 and specific axial resistance of *Ri* = 100Ω*cm*. **B**, Similar calculation for the NeuroMorpho.Org database as Figure 4A but with appended somata that do not receive synaptic inputs (see Methods for more details). Here, colours indicate the radius of the appended soma.

**Fig S4.**
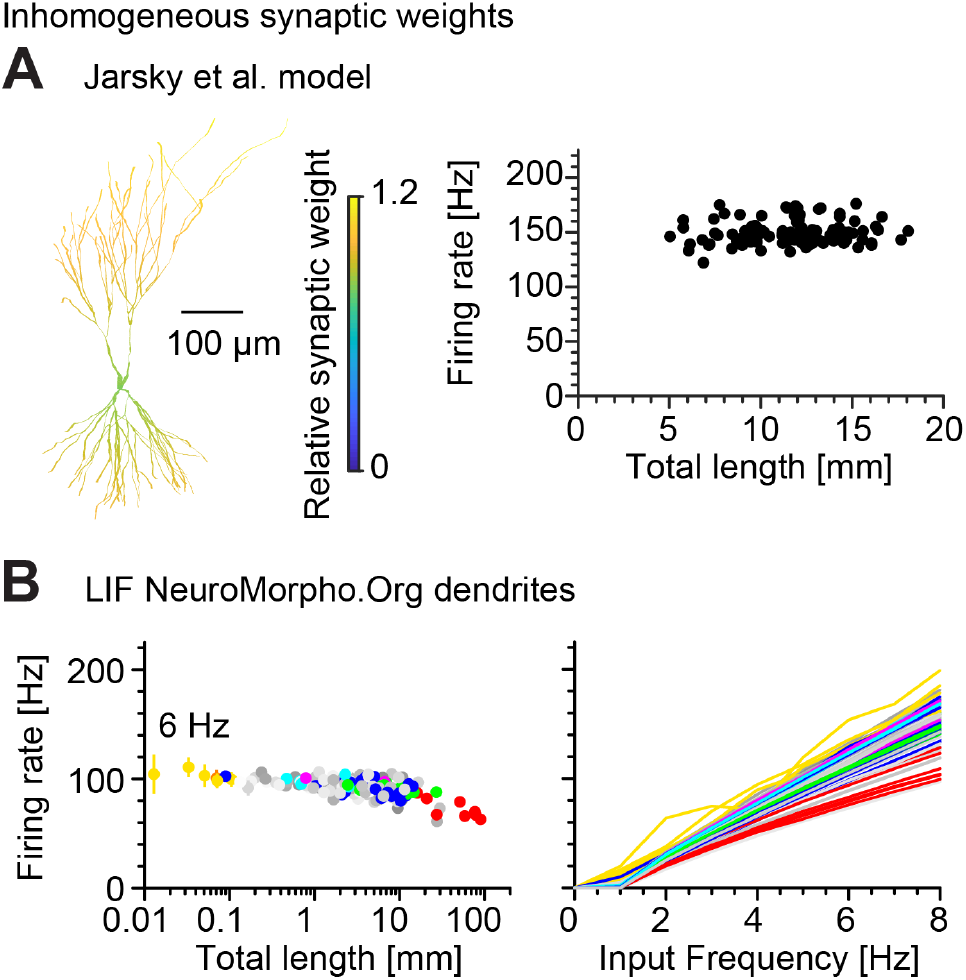
Effect of inhomogeneous distance-dependent synapse weights on dendritic constancy. In these plots, synapses were scaled from 0.8× to 1.2× in a linear relation with distance (path length) from soma. **A**, Analogous to Figure 6B, the model by Jarsky et al. (2005) with inhomogeneous synapse weights. **B**, Analogous to Figure 8A, the LIF model in NeuroMorpho.Org morphologies with inhomogeneous synapse weights.

**Fig S5.**
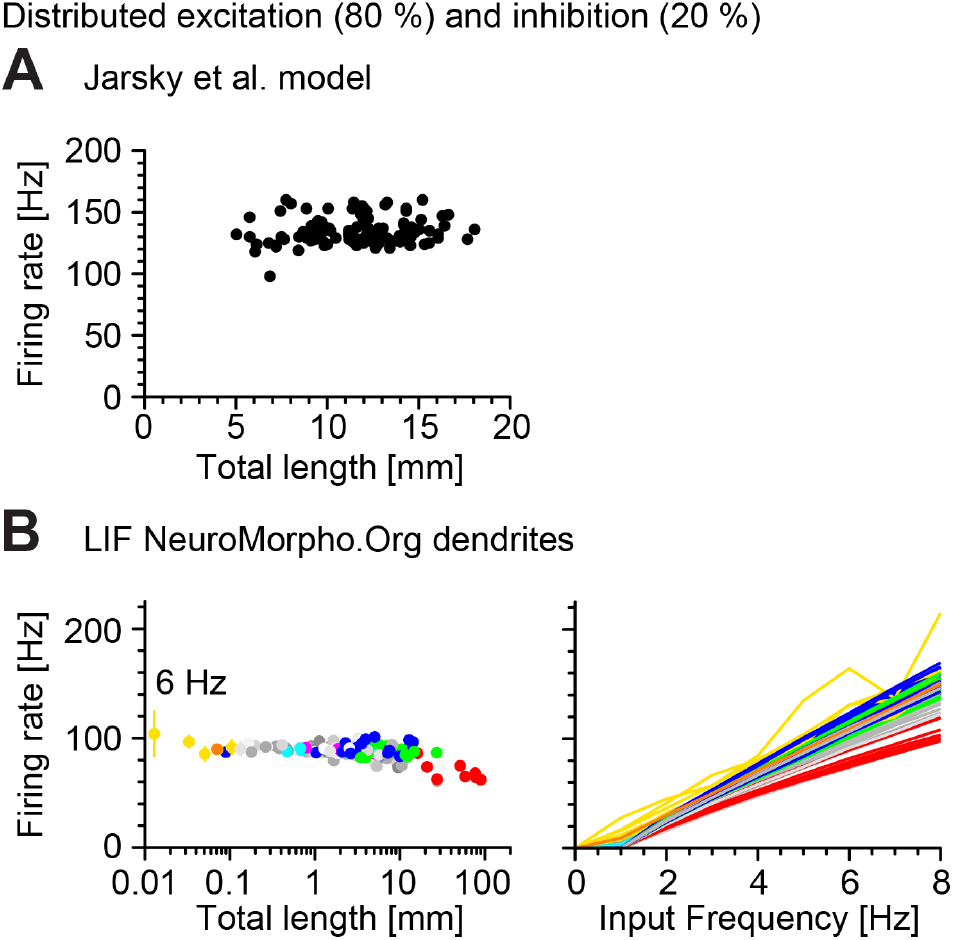
Effect of inhibitory synapses on dendritic constancy. In these plots, 20% of all synapses were randomly selected to have a reversal potential of −80*mV*, which renders them inhibitory synapses. **A**, Analogous to Figure 6B, the model by Jarsky et al. (2005) with 20% inhibitory synapses. **B**, Analogous to Figure 8A, the LIF model in NeuroMorpho.Org morphologies with 20% inhibitory synapses.

**Fig S6.**
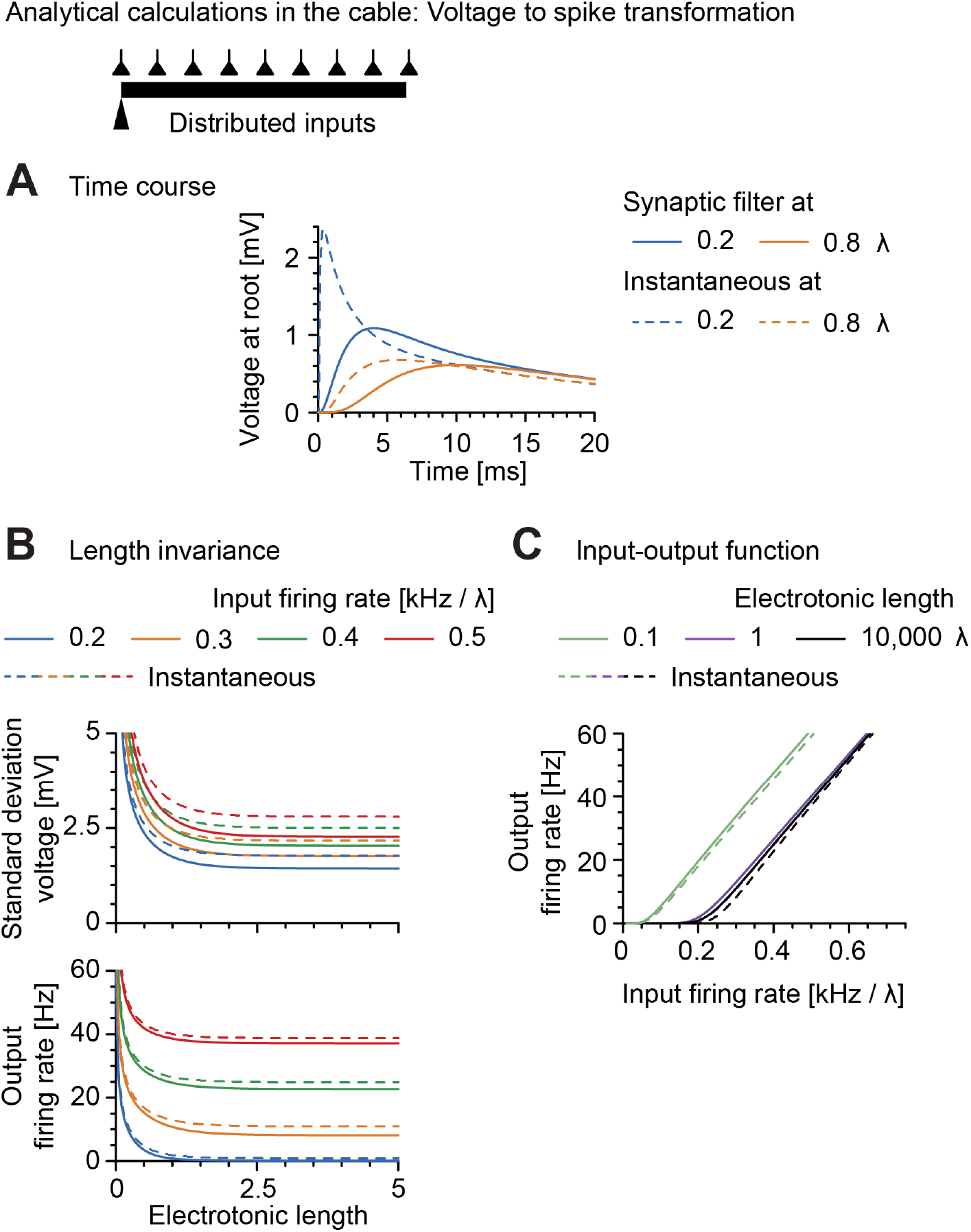
Transformation of voltage fluctuations into spikes. **A**, Voltage response as a function of time at the proximal end of a sealed dendrite of electrotonic length 1 to an instantaneous (dashed lines) and synaptically filtered (double exponential *τ*_*rise*_ = 0.5*ms*, *τ*_*decay*_ = 2.5*ms*, solid lines) current injection with magnitude *a* = 1. Blue lines show responses at 0.2 and yellow lines at 0.8 electrotonic distance. The time constant of the membrane was *τ* = 20*ms*. **B**, Top panel, subthreshold proximal voltage variance as a function of electrotonic length for different input rates per unit electrotonic length: 200, 300, 400, and 500*Hz*. *a* = 2.5 and *τ* = 20*ms*. Bottom panel, firing rate as a function of electrotonic length for different input rates per unit electrotonic length: 200, 300, 400, and 500*Hz*. *a* = 2.5 and *τ* = 20*ms*. As in **A**, dashed lines for instantaneous and solid lines for filtered current injections. **C**, Output firing rate as a function of afferent rate for dendrites of different electrotonic lengths: 0.1, 1, and 10, 000. *a* = 2.5 and *τ* = 20*ms*. As in **A**, dashed lines for instantaneous and solid lines for filtered current injections.

**Table S1.**
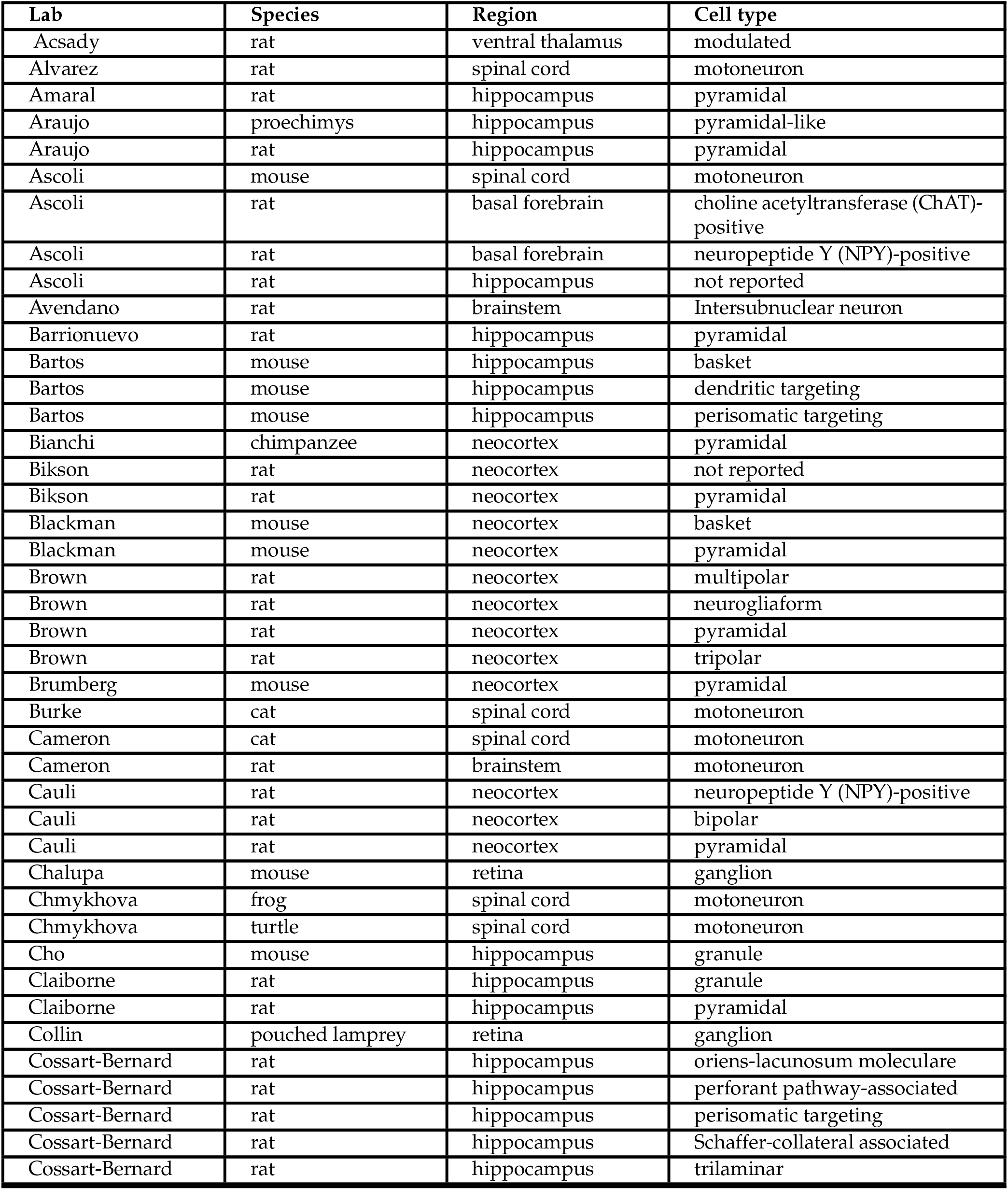
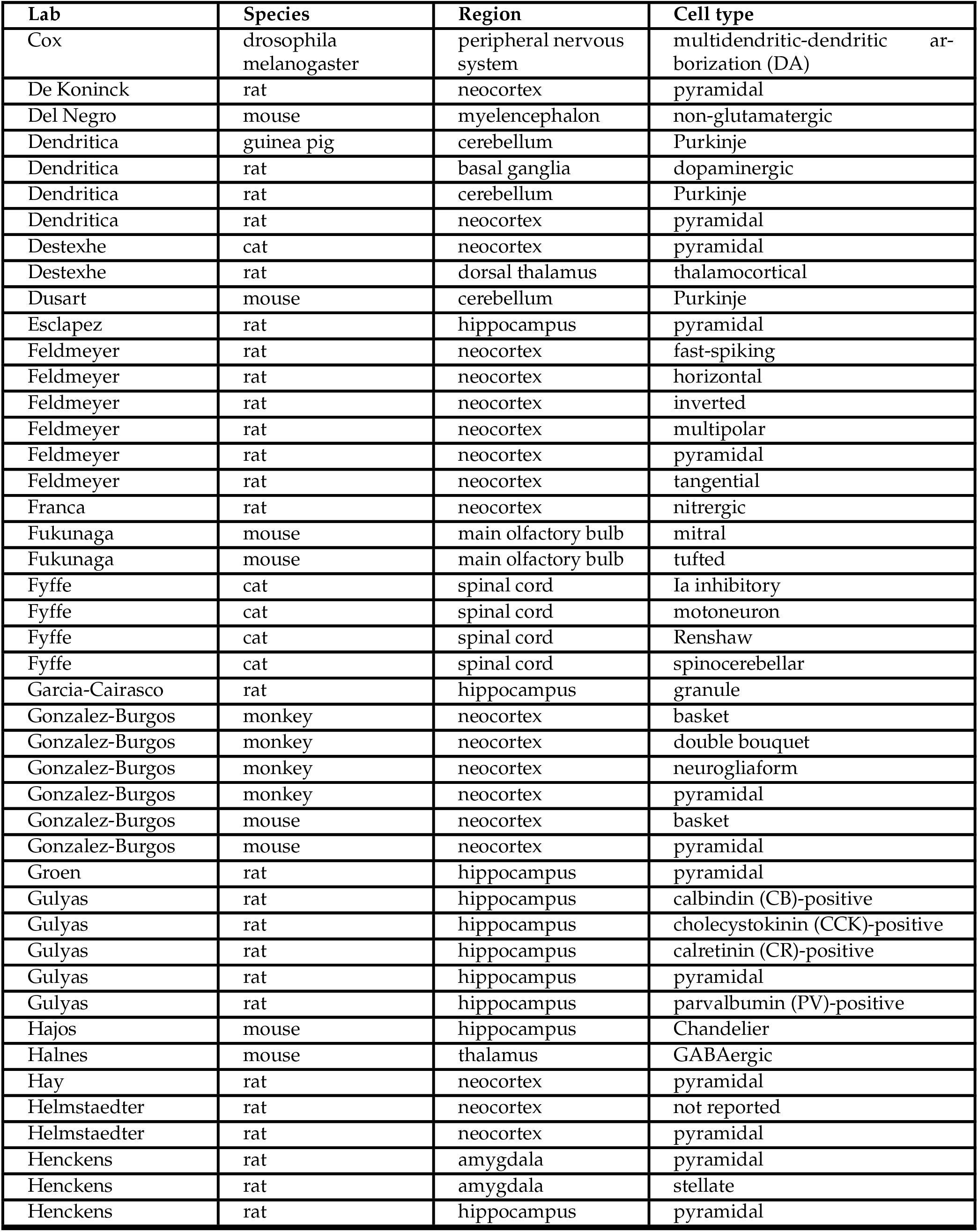
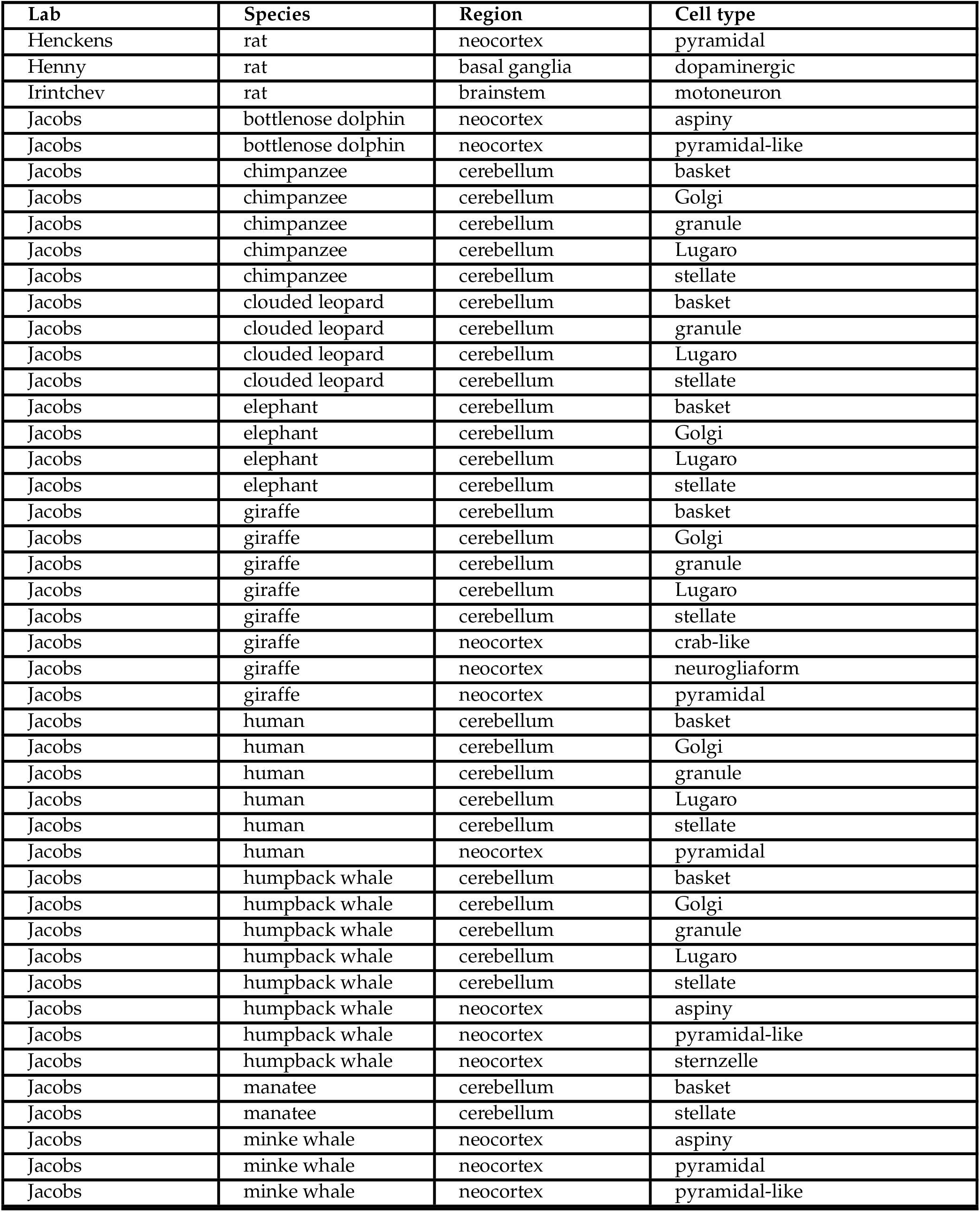
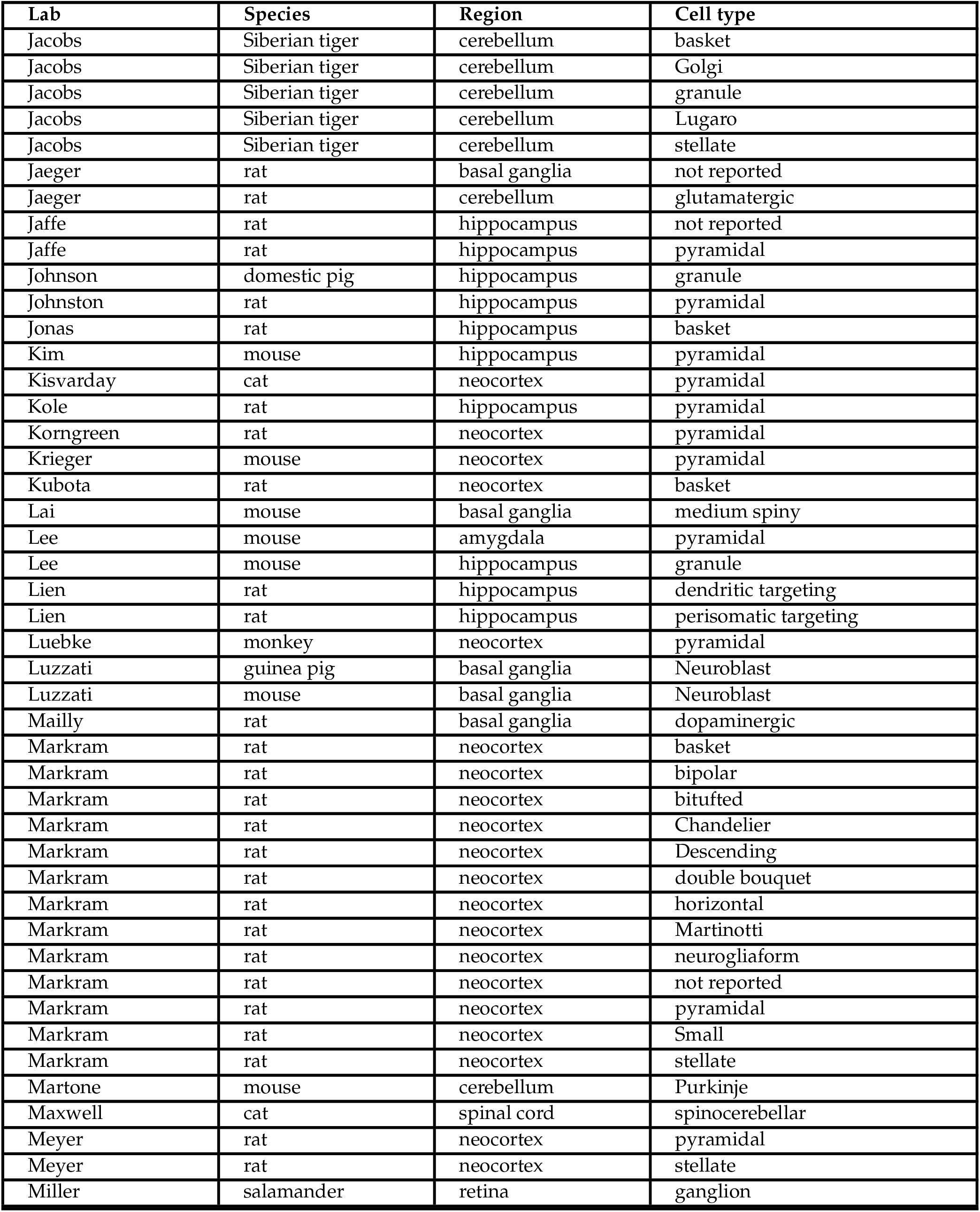
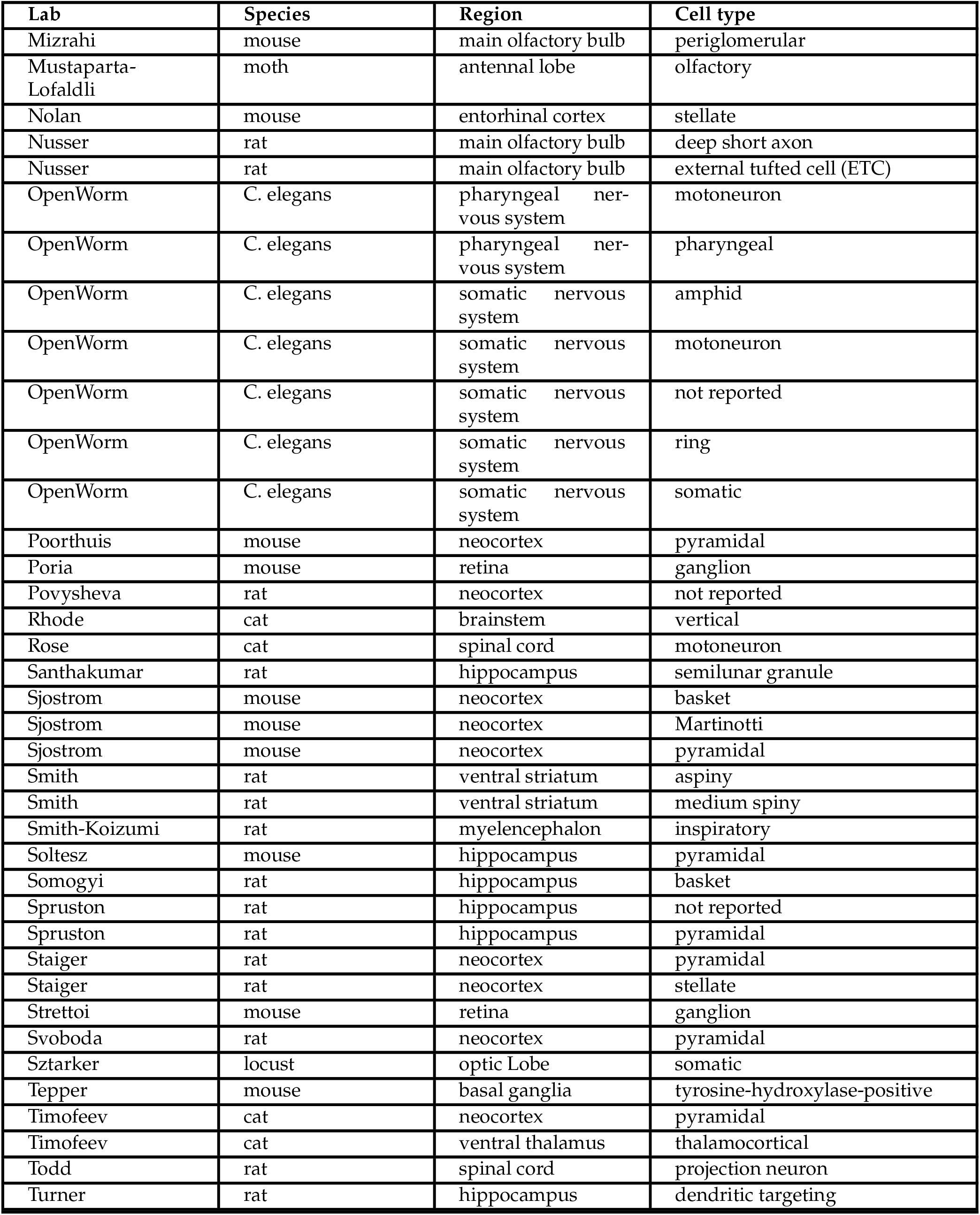
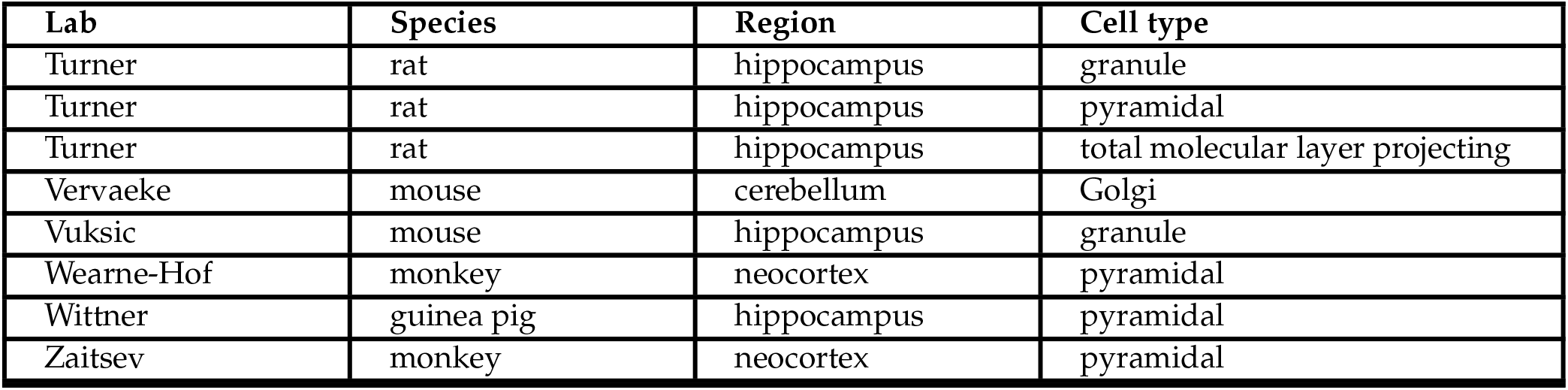
Selected datasets from NeuroMorpho.Org

